# Sequential estimation of nonstationary oscillator states and stimulus responses

**DOI:** 10.1101/2024.06.24.600503

**Authors:** Joseph N. Nelson, Alik S. Widge, Théoden I. Netoff

## Abstract

Neural oscillations have been linked to multiple behaviors and neuropsychiatric disorders. The individual contributions to behavior from both oscillations and non-oscillatory activity are still unclear, complicating efforts to link neurophysiology to cognition, hindering the discovery of novel biomarkers, and preventing the development of effective therapeutics. To overcome these hurdles, it will be critical to investigate the biological origins of neural oscillations by characterizing the dynamic properties of different brain regions. The dynamical regime for a population of neurons generating oscillations in neural recordings can be discovered by stimulating the population and recording its subsequent response to stimulation. There are different dynamical regimes that can produce population-level neural oscillations. For certain dynamical regimes, like that of a nonlinear oscillator, the phase response curve (PRC) can help differentiate the dynamic state of the population. The PRC can be measured by stimulating the population across different phases of its oscillatory state. However, neural dynamics are non-stationary, so neural oscillations will vary in frequency and amplitude across a recording and the PRC can change over time. This non-stationarity could bias a PRC estimated from an electrophysiological experiment, preventing accurate characterization of a neural population’s dynamics. This necessitates tools that can operate online to trigger stimulation and update PRC estimates. To that end, we develop online methods for tracking non-stationary oscillations and recovering PRCs corrupted by estimation errors. We validate the performance of our non-stationary oscillation estimator compared to both a known ground truth model and an alternative phase estimation approach. We demonstrate that a PRC can be recovered online under different random error conditions *in silico* and that a similar amplitude response curve (ARC) can be estimated from physiologic data using online methods compared to offline approaches.

## 1 Introduction

Neural oscillations are a ubiquitous feature in brain recordings, such as local field potentials (LFP) and electroencelphalograpy (EEG). Neural oscillations are categorized into specific frequency ranges, such as theta (4-8 Hz), alpha (8-12 Hz), and beta (12-30 Hz), that each show different changes during motor movements, sensory processing, and cognitive tasks (Canolty and Knight 2010). For instance, changes across these bands can decode the level of depressive symptoms being experienced by epilepsy patients undergoing intracranial EEG (Scangos et al. 2021). Enhanced beta power in the basal ganglia correlates with bradykinesia and rigidity for some Parkinson’s disease patients (Kehnemouyi et al. 2021). For certain psychiatric disorders, like addiction and post-traumatic stress disorder, there are inconsistent changes in spectral power across research studies, with conflicting trends in band-specific power reported (Newson and Thiagarajan 2019). Further, non-oscillatory aperiodic neural activity, with 1*/f* -like spectral features, may be a better classifier for schizophrenia than frequency-band specific features (Peterson et al. 2023). These inconsistencies in the clinical relevance of neural oscillations across different neurologic and psychiatric disorders indicate a need to better characterize the biological origins of these oscillations. Understanding the biological mechanism which generate oscillations is also important for neuromodulation, because stimulation can have different effects depending on an oscillation’s state. For example, electrical stimulation at rising or falling phases of a beta rhythm induced opposite changes in synaptic connectivity (Zanos et al. 2018). The results of this study were replicated in a model network of neurons with plastic synapses that received a beta oscillatory input, rather than generating its own beta oscillations (Shupe and Fetz 2021), which leaves the question of the neural oscillations’ origins open.

The biophysical mechanisms that generate neural oscillations are still debated (Doelling and Assaneo 2021). Some theories posit that oscillations are signals from populations of neurons behaving collectively as nonlinear oscillators with stable **limit cycles** (Weerasinghe et al. 2021; Doelling and Assaneo 2021). These theories have been expanded to provide a hypothetical mechanism for inter-areal brain communication using the theory of weakly coupled oscillators - each neural population is a nonlinear oscillator and these populations interact through exchanged perturbations in their phases and amplitudes. However, many areas of the cerebrum are dynamically in an asynchronous state, a state in which the mean spiking correlations between neurons are low and neurons’ spike trains are irregular (Brunel 2000; Renart et al. 2010), rather than in an oscillatory state. This asynchronous state is stabilized in the cortex through a balance of excitatory and inhibitory synaptic currents (Shadlen and Newsome 1998; Vreeswijk and Sompolinsky 1996). In the mean-field theory of recurrent excitatory and inhibitory spiking networks, this balanced asynchronous state is modeled as a dynamical system with a stable fixed point (Gerstner et al. 2014). A system with a stable fixed point will not respond to perturbations in the same ways as a nonlinear oscillator with a stable limit cycle (Strogatz 2014). In the asynchronous state, finite populations of irregular spiking neurons can respond to rapid inputs with damped oscillations (Schwalger and Chizhov 2019), display fluctuations in population-level activity due to their finite size (Gerstner et al. 2014), and feature peaks in the power spectral density of their spike trains in common neural oscillation frequency bands, such as gamma (30-50 Hz) (Richardson 2008). This alternative mechanism to limit cycles for oscillation generation, called **quasi-cycles**, in finite-size populations has been studied in neuroscience (Wallace et al. 2011), ecology (A. J. McKane and Newman 2005), and epidemiology (Alonso, Alan J McKane, and Pascual 2006).

The limit cycle and quasi-cycle mechanisms for oscillation generation are not easily distinguished using the power spectral density of recorded neural oscillations (Wallace et al. 2011). Instead, the two mechanisms can be distinguished based on the response of a population of neurons to perturbations, such as electric and optogenetic stimulation. For example, nonlinear oscillators can be reduced to one-dimensional phase models with a phase response curve (PRC), which quantifies the effect of an external perturbation on the phase of a limit cycle (Ermentrout and Terman 2010). Delivering stimulation at different phases of an oscillation (phase-locked stimulation) is one method of estimating an oscillator’s PRC, by measuring how the oscillator’s phase changes in response to the applied stimulation. The particular form of the estimated PRC will reveal the type of oscillator - a strictly positive PRC with only phase advances corresponds with a Type-1 oscillator, while a PRC that is both positive and negative with phase advances and delays implies a Type-2 oscillator (Ermentrout and Terman 2010). If the state of the system were static, we could collect these stimulus-induced phase changes, estimate the system’s PRC, and use this estimated PRC throughout an electrophysiological experiment to controllably shift the phase of the neural population oscillator. However, in the brain, behavioral factors such as learning and attention can change the operating point, or state, of a population of neurons (Huang 2021). A change in state of a neural population can alter the dynamical properties of the system, such as the population’s response to an external input. A population oscillator could transition into the asynchronous state in which a PRC is undefined; or transition from one oscillatory regime to another (e.g., Type 1 to Type 2). As a further complication, a network of neurons in an asynchronous state can even transition into a global oscillatory state, while the spike trains of individual neurons remains irregular (Brunel 2000). These changes in state of the neural population’s dynamics would therefore necessitate an update to our PRC estimate.

There is a need for methods that can adaptively estimate a PRC “on-the-fly” as new information is obtained about the system through experimental perturbations.

In this study, we discover a PRC using Bayesian inference (Garnett 2023) to refine our estimate under uncertainty as we receive new information from an online perturbation experiment. Bayesian inference has had great success in biology and biomedical engineering, from simulation-based inference of neuroscience models using experimental data (Gonçalves et al. 2020) to parameter optimization of wearable devices (Kim et al. 2017). In neuromodulation, deep brain stimulation parameters have been efficiently optimized using Bayesian optimization, a sequential design strategy that performs global optimization on an objective function (Connolly et al. 2021; Grado, Johnson, and Netoff 2018; Nagrale et al. 2023). The statistical model of this typically unknown objective function is a Gaussian process, a stochastic process over random variables jointly distributed as a multivariate normal distribution (Rasmussen and Williams 2005; Garnett 2023), where the random variables are experimental measurements. This model can be used for regression tasks for which the form of the underlying function is unknown, which is typically the case for the PRC of an unknown oscillatory system. We demonstrate how a PRC estimate can be updated as different phases are sampled using Gaussian process regression, an approach we refer to as Adaptive Oscillation Response Estimation (AORE).

To deliver phase-locked stimulation to the brain in order to estimate a PRC, the phase of a neural oscillation must be estimated online to trigger the stimulating device. Neural oscillations commonly vary in frequency, amplitude, and mean over time (Donoghue, Schaworonkow, and Voytek 2022) and these oscillations are recorded from electrodes that will corrupt the signal with measurement noise. An effective phase estimator must be able to track these non-stationary oscillations and filter the measurement noise to avoid large estimation bias. Neural oscillations are typically processed with a bandpass filter and the Hilbert transform (the so called filter-Hilbert method), which has been a useful approach for delivering phase-locked stimulation in the nervous system (Zrenner et al. 2020; Cagnan et al. 2017; Schatza et al. 2022). However, this filter-Hilbert approach to oscillation estimation can lead to edge artifacts in the estimated phase without careful tuning of the filter window. The time resolution of the filter-Hilbert is also limited by the chosen window length of the filter. A model-based oscillation estimation approach would avoid window edge artifacts, allow more flexibility in the time resolution of the phase estimate, and allow researchers to instantiate their hypotheses of the neural oscillation features (like non-sinusoidal waveforms and stochastic walks in the center frequency) into the estimator’s model. There have been a variety of approaches to estimate the phase of neural oscillations, including combining the filter-Hilbert with an autoregressive model to compensate for filter phase delays (Schatza et al. 2022). Other model-based phase estimation approaches exist, such as the state-space phase estimator (SSPE) (Wodeyar et al. 2021), which has been shown to track multiple co-existing oscillations in silico and neural oscillations from EEG and LFP. The SSPE uses a Kalman filter for sequential state estimation of oscillations, with a damped harmonic oscillator model in the filter. However, the SSPE does not explicitly include the frequency of the oscillation as a state variable. Including a frequency variable could be useful for tracking the frequency drift of a neural oscillation. Further, the SSPE uses a linear model, which does not have a limit cycle (Strogatz 2014). By using a nonlinear model within a Kalman oscillation estimator, an investigator could differentiate between self-sustaining oscillation epochs and damped oscillatory periods using a bifurcation parameter. Therefore, we developed an unbiased, nonlinear sequential state estimator of non-stationary oscillations, which we refer to as Adaptive Oscillation State Estimation (AOSE), to track non-stationary oscillations in a ground truth model test and from physiological oscillations (Cagnan et al. 2017). We then use the AOSE oscillation tracker with AORE to adaptively estimate PRCs from both a computational model oscillator (Jansen and Rit 1995) and a previous phase-locked physiological experiment (Cagnan et al. 2017). We believe this adaptive framework for estimating both the oscillation and the oscillatory stimulus-response will benefit future studies investigating the dynamical regimes of different brain regions.

## 2 Methods

The Adaptive Oscillation State Estimator (AOSE) performs sequential state estimation of non-stationary oscillations using an Extended Kalman Filter (section 2.3). We check the consistency of the AOSE in recovering an unbiased state estimate with an accurate estimate of the error statistics using a Monte Carlo truth model test (TMT) and using the Normalized Estimation Error Squared (NEES) statistic (section 2.4). Population-level neural dynamics are commonly state-dependent, such that the population’s response to stimulation may vary as the biological system transitions into different dynamical regimes (e.g., a transition from a Type 1 to a Type 2 oscillator). This dynamic transition would then change the shape of the oscillatory system’s phase response curve (PRC). The Adaptive Oscillation Response Estimator (AORE) combines the AOSE with Gaussian Process Regression to discover the biological dynamical system’s PRC as data is recovered from a sequential sampling experiment (section 2.5). We deliver synthetic phase-locked stimulation to the Jansen-Rit model to extract the system’s PRC (section 2.1), which is used to test the performance of the AORE. Accelerometry data recorded from Essential Tremor patients’ tremulous hands in a previous study (Cagnan et al. 2017) is analyzed offline and online (section 2.2) to test AOSE on a physiologic dataset with a known phase-dependent amplitude response curve (ARC).

### 2.1. Simulating a non-stationary neural oscillator with the Jansen-Rit model

To model neural oscillations from LFP and EEG, we used the Jansen-Rit model (Jansen and Rit 1995) that includes one inhibitory and two excitatory neural populations, with second-order differential equations that model the population average membrane potential dynamics. The Jansen-Rit model includes six ordinary differential equations:

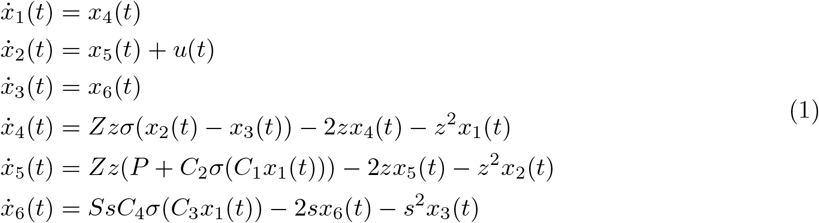

where *Z, S*, and *s* are constants which determine the maximum amplitude of the synaptic activity, *C*_*n*_ referes to the coupling constants between populations and P refers to an external stimulus to the main pyramidal neural population from the interneuron populations. The membrane potential is transformed into a firing rate using the following sigmoid function

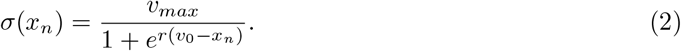

The parameter *v*_*max*_ is the maximum firing rate, *v*_0_ is the value for which a rate of 50% is achieved and *r* corresponds to the slope. We set *Z* = 3.25*mV*, *S* = 22*mV*, *z* = 100*s*^−1^, *s* = 50*s*^*−*1^, *v*_0_ = 6*mV*, *v*_*max*_ = 5*s*^*−*1^, and *r* = 0.56*mV* ^*−*1^, *C*_1_ = *C, C*_2_ = 0.8*C, C*_3_ = 0.25*C, C*_4_ = 0.25*C, C* = 135 (Grimbert and Faugeras 2006).

The Jansen-Rit model was simulated as stochastic differential equations using a Wiener process

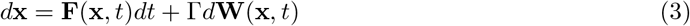

where **W** denotes a Wiener process and Γ∈ ℝ^6*×*6^ is a process noise covariance matrix. The Jansen-Rit model was simulated using Diffrax, a library of numerical differential equation solvers (Kidger 2022). We used the Euler Maruyama solver with Diffrax’s Virtual Brownian Tree class to simulate the Wiener process, and a constant step size was used. The main pyramidal population of the Jansen-Rit model was stimulated using a monophasic pulse with width 0.01 seconds and amplitude of 50. The monophasic pulse was linearly interpolated to the solver’s required timescale. To deliver a phase-locked pulse to the model, the Adaptive Oscillation State Estimator was used to estimate the oscillator state at every time step (Figure 1). For each trial, the oscillator phase shift from stimulation was computed. The phase shift was calculated as the phase difference between the stimulated model and the unperturbed model (with the same initial condition) 1 second after stimulation was delivered. To speed up the simulation time, the simulation pipeline was just-in-time (jit) compiled in JAX. The phase difference was computed as the angle with minimum arc length between two points on the unit circle (Abuzaid, Mohamed, and Hussin 2012):

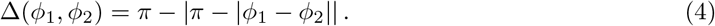

**Figure 1:**
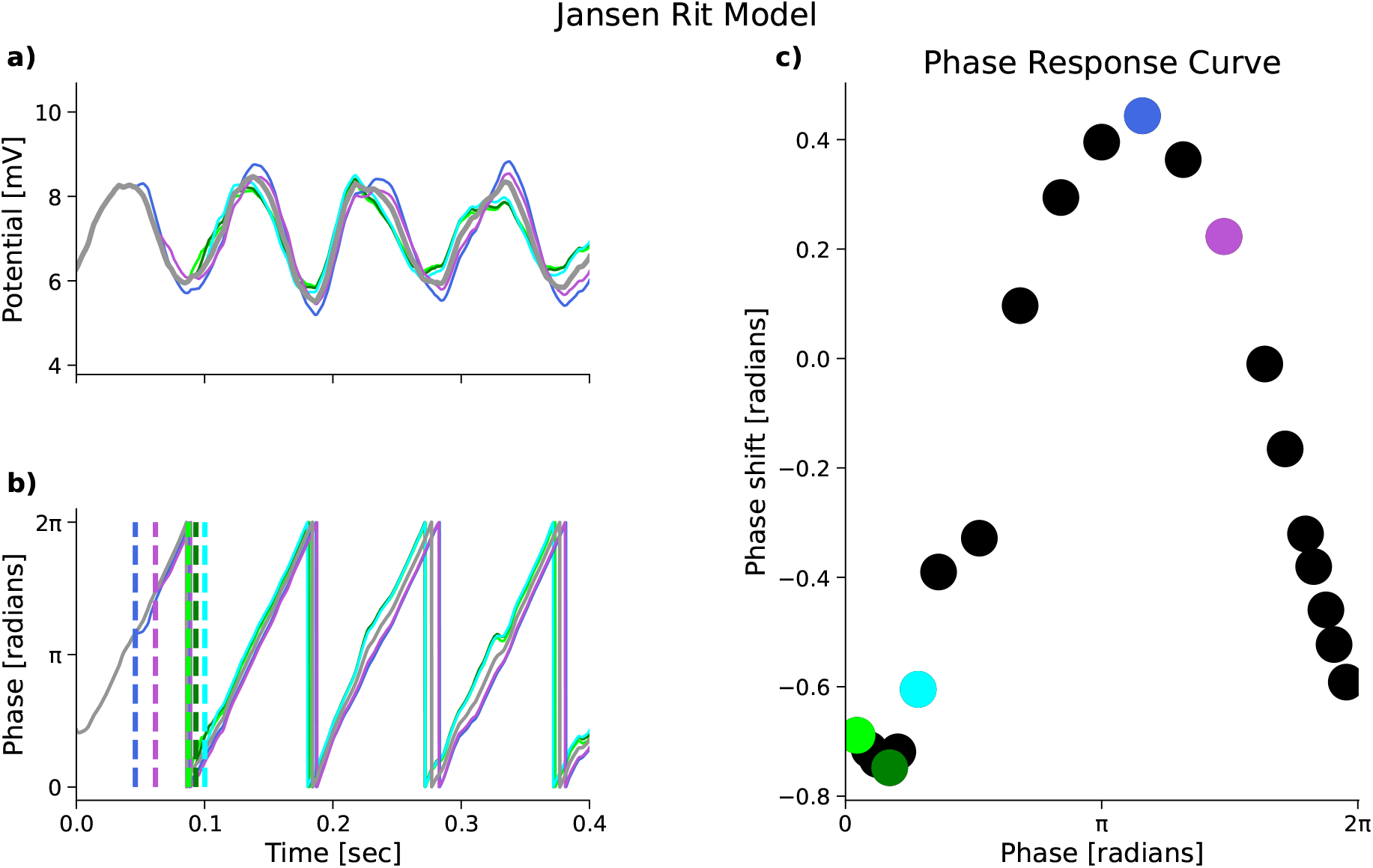
Single pulse phase-locked stimulation delivered to the second state variable of the stochastic Jansen-Rit (JR) model recovers the oscillator’s phase response curve (PRC) (the direct method). For each simulation, a single phase-locked monophasic pulse was delivered to the model’s main pyramidal neural population. The unperturbed simulation is color coded in gray and the phase-locked simulations are in different cool colors (lime, green, aqua, royal blue, and medium orchid). (a) JR pyramidal population membrane potential. Stimulation delivered at different phases of the neural oscillator result in both phase delays and phase advances. (b) Phase of JR oscillations, estimated using the Adaptive Oscillation State Estimator (AOSE). Phase-locked stimulation times are denoted with dashed lines, with the same color code as the phase of stimulation. (c) Points near the JR PRC sampled using the direct method (dots). Phase shifts in (a) and (b) are visualized as shaded dots, with the same color code. The JR PRC, with the chosen parameters, is a Type 2 PRC with both phase delays and phase advances (Ermentrout and Terman 2010).

### 2.2. Signal processing of experimentally observed physiologic oscillations and amplitude response curves

In a previous study (Cagnan et al. 2017), researchers suppressed the hand tremors of essential tremor (ET) patients by delivering deep brain stimulation (DBS) to the ventrolateral thalamus, phase-locked to the ET patients’ hand tremor rhythm. This tremor rhythm typically oscillates around 4-8 Hz (Figure 2b). They found that stimulating the thalamus at certain tremor phases resulted in significant changes to the amplitude of the tremor rhythms. Their analysis used an offline signal processing pipeline. Briefly, Cagnan and colleagues used a zero-phase bandpass filter to extract the tremor band rhythms and computed the instantaneous phase and amplitude of these rhythms using the Hilbert Transform. The DBS-induced change in tremor amplitude was computed by segmenting the data offline into trial epochs consisting of a 1-second pre-stimulus interval and a 5-second interval of phase-locked stimulation; they then took the difference between the average tremor amplitude in the last second of the stimulation interval and the average amplitude of the pre-stimulus interval, and normalized this difference by the pre-stimulus average. We made a similar offline signal processing pipeline in MNE-python (Gramfort et al. 2013) with a filter length of 1-second, low and high bandpass limits *±*2 Hz about the frequency at the peak of pre-stimulus power spectral density in the tremor band (4-8 Hz), a lower passband of 2 Hz and an upper passband of 5 Hz. The bandpass filter used a Hamming window. The data was downsampled by a factor of 10 using MNE’s resample method, which includes an anti-aliasing filter. The power spectral density was estimated using Welch’s method, with a 0.5 second Hamming window and 1/2 window overlap. For participants 1, 3, 4L, 5, and 6 (labels for ET participants in Cagnan et al. 2017), the average targeted tremor phase and average tremor amplitude change for every stimulation block were compiled into a feature set, which we used to calculate the offline PRC estimates. We used the ordinary least squares method from the statsmodels package to fit linear models to the particpants’ feature sets. We chose a collection of 15 different linear models, each with a different set of basis functions, to test on each feature set (Table 1). We computed the Bayesian Information Criterion (BIC) using statsmodels for each linear model on these participants’ feature sets. We selected the model with the lowest BIC score as the offline PRC for each participant (highlighted in bold red in Table 1). For participant 1, two of the linear models both resulted in the same lowest BIC score (BIC = *−*1092); we chose the simpler model with a constant coefficient (1), rather than a linear term (*x*).

**Table 1:** Offline PRC model selection.

| Basis | BIC |  |  |  |  |
| --- | --- | --- | --- | --- | --- |
|  | 1 | 3 | 4L | 5 | 6 |
| $x, \sin(x), \cos(x), \sin(2x), \cos(2x), \sin(3x), \cos(3x)$ | -1044 | -1053 | -1044 | -704.5 | -827.2 |
| $x, \sin(x), \cos(x), \sin(2x), \cos(2x), \sin(3x)$ | -1078 | -1057 | -1048 | -704.6 | -830.8 |
| $x, \sin(x), \cos(x), \sin(2x), \cos(2x)$ | -1083 | -1061 | -1052 | -708.7 | -831.2 |
| $x, \sin(x), \cos(x), \sin(2x)$ | -1088 | -1065 | **-1053** | -712.5 | -834.5 |
| $x, \sin(x), \cos(x)$ | **-1092** | -1070 | -1047 | -710.0 | -837.2 |
| $x, \sin(x), \cos(x), \cos(3x)$ | -1088 | -1065 | -1042 | -709.1 | -834.3 |
| $x, \sin(x), \cos(x), \cos(3x), \sin(3x)$ | -1083 | -1061 | -1039 | -706.0 | -834.3 |
| $x, \sin(x), \cos(x), \cos(3x), \cos(2x), \sin(3x)$ | -1079 | -1057 | -1039 | -702.3 | -830.5 |
| $1, \sin(x)$ | -1088 | -1073 | -1050 | -668.7 | -834.8 |
| $1, \sin(x), \cos(x)$ | **-1092** | -1070 | -1047 | -707.8 | **-839.4** |
| $1, \sin(x), \cos(x), \sin(2x)$ | -1087 | -1066 | -1051 | **-713.1** | -835.3 |
| $1, \sin(x), \cos(x), \sin(2x), \cos(2x)$ | -1083 | -1061 | -1051 | -709.2 | -831.8 |
| $1, \sin(x), \cos(x), \sin(2x), \cos(2x), \sin(3x)$ | -1078 | -1058 | -1047 | -704.8 | -829.8 |
| $1, \sin(x), \cos(x), \sin(2x), \cos(2x), \sin(3x), \cos(3x)$ | -1074 | -1053 | -1043 | -705.0 | -826.3 |
| $1$ | -1090 | **-1077** | -1045 | -668.3 | -838.6 |

**Figure 2:**
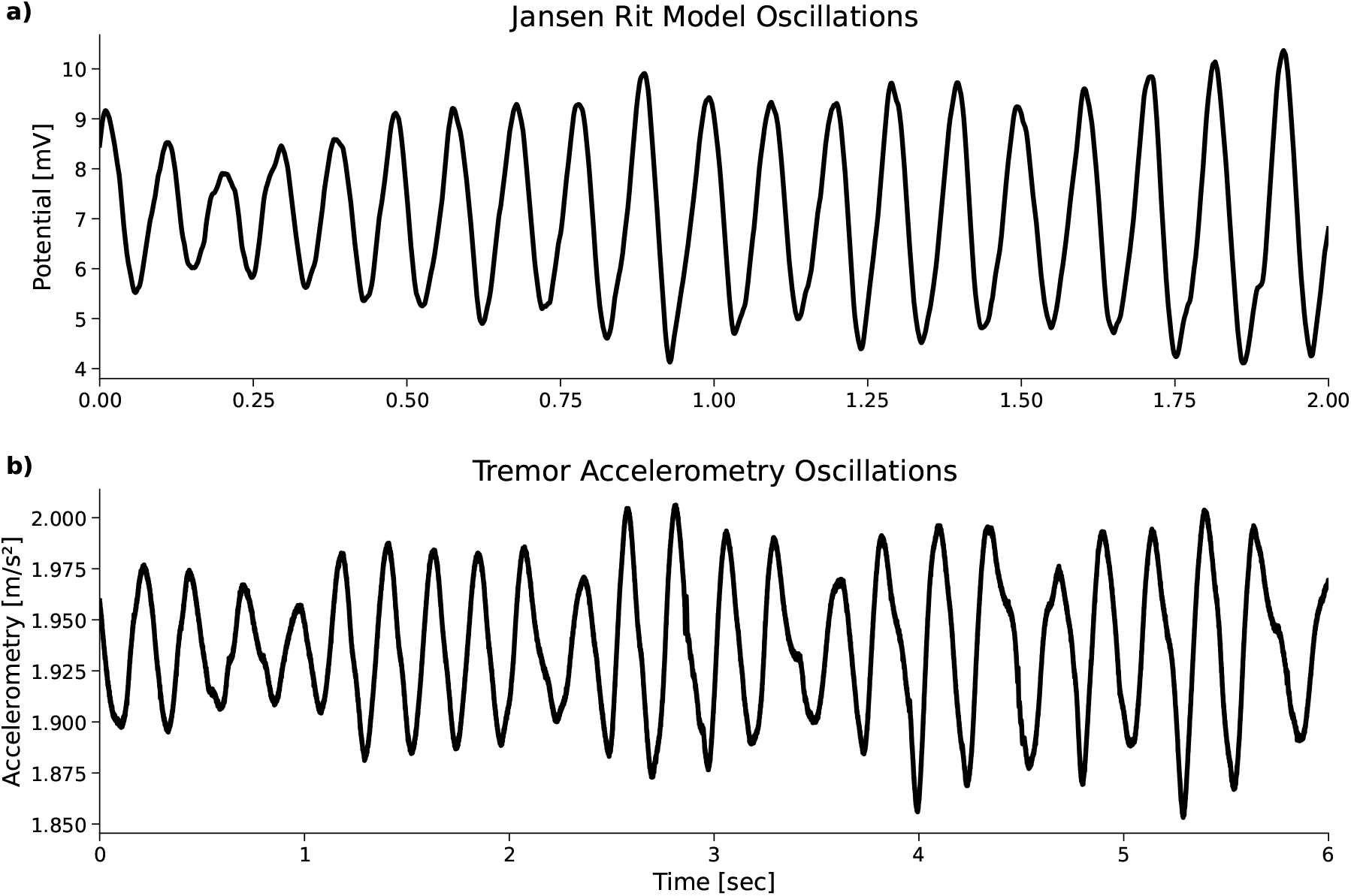
Oscillations in a computational model and peripheral muscle tremors. Both examples are characterized by sequences of decaying and sustained oscillations and distortions in the shape of their waveforms. (a) Membrane potential of the main pyramidal population in the Jansen-Rit (JR) model. With our chosen parameters from (Grimbert and Faugeras 2006), the JR oscillation center frequency was approximately 10 Hz. (b) Hand tremors from Participant 5 in Cagnan et al. 2017. Tremors were sensed using an accelerometer attached to the participant’s tremulous hand. Tremor oscillations center frequency was around 4-6 Hz across participants.

In this study, for the online estimation approach, we estimated the change in the average amplitude by storing the pre-stimulation (PRE) and post-stimulation (POST) accelerometry amplitude estimates from the online oscillation state estimator into first-in-first-out (FIFO) queues. The queue data structures were set to only store, at most, 2 seconds of data. The PRE queue stored data one second before the stimulus block began and the POST queue stored the last one second of data during the stimulus block. The PRE and POST queue data structures included a class boolean to indicate whether the streaming data was currently in the pre-stimulus block (for PRE queue) or stimulation block (for POST queue), which was set using the stimulation trigger times saved with the Cagnan dataset. Once a 1/2 second passed with no stimulus trigger events, the collected amplitude values from the POST queue were averaged. Once the next 1-second stimulation block began, the amplitude values from the 1-second period between stimulation blocks stored in the PRE queue were averaged. The average target phase was estimated using another FIFO queue (denoted as the PHASE queue). The PHASE queue kept a boolean class variable to indicate whether the streaming accelerometry data was currently in the stimulation block.

### 2.3. Sequential state estimation of non-stationary oscillations

The Adaptive Oscillation State Estimator utilizes an Extended Kalman Filter to track non-stationary oscillations. The Extended Kalman Filter (EKF) is a sequential state estimation algorithm that uses a nonlinear model to predict the state(s) of a dynamic system, and updates the state estimates using measurements assumed to be corrupted by additive white Gaussian noise. An EKF assumes the underlying dynamic system to be estimated is

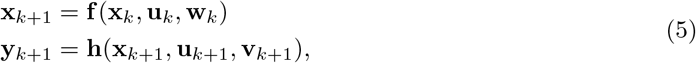

where **x**_*k*_ ∈ ℝ^*n*^ is the dynamic system’s vector of state variables, **u**_*k*_ ∈ ℝ^*m*^ is an externally applied input to the system, **w**_*k*_ ∼ *N* (0, *Q*) is the process with covariance noise *Q* ∈ ℝ^*n×n*^ and **v**_*k*_ ∼ *N* (0, *R*) is the measurement noise with covariance *R* ∈ ℝ^*m×m*^. There are *n* state variables and *m* input variables. **f** is the state dynamics function, a nonlinear discrete-time dynamic system, and **h** is the sensor (or measurement) function, a vector of potentially nonlinear algebraic equations. The EKF has a state prediction step and a measurement correction step as follows:

#### Prediction

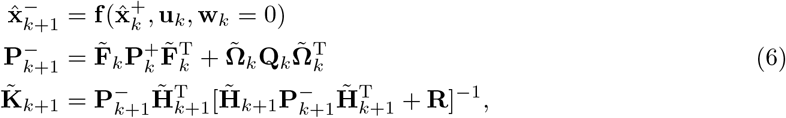

#### Correction

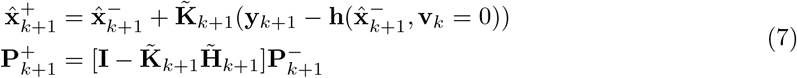

For every time step *k* ∈ ℤ_≥0_, 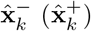 is the state estimate from the state prediction (measurement update) step. 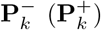 is the estimation error covariance matrix from the state prediction (measurement update) step. The state transition matrix 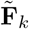, which is used by a *linear* (rather than nonlinear, like the EKF) Kalman Filter to propagate the state estimate forward in time, is related to the Jacobian of the state dynamics function, 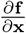, as

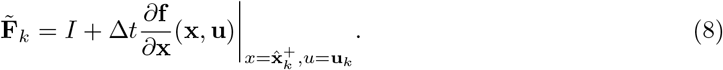

The sensor matrix, which maps states to measurements, 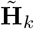 is related to the Jacobian of the sensor function, 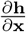, as

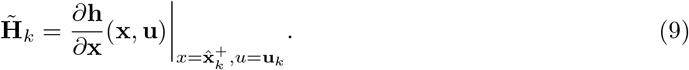

**y**_*k*_ ℝ^*p*^ is the data from measurements and 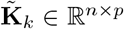 is the Kalman gain matrix that optimally blends the measurements and state estimates.

The nonlinear model for the EKF is an extension of the Stuart-Landau model (Kuznetsov 2004) with additional appended states to allow for non-stationary oscillation frequency, mean, and amplitude. The Stuart-Landau model can generate oscillations through a Hopf bifurcation (Kuznetsov 2004). The extended Stuart-Landau model is represented as

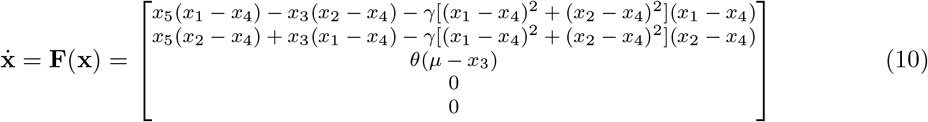

with state vector *x* = [*x*_1_, *x*_2_, *x*_3_, *x*_4_, *x*_5_]^*T*^. Variables *x*_1_ and *x*_2_ are the states of the original Stuart-Landau model oscillator, *x*_3_ is the mean-reverting frequency, *x*_4_ is the mean that the Stuart-Landau model oscillates about, and *x*_5_ is the bifurcation free-parameter of the oscillator model. *γ* is a coupling parameter that weights the nonlinear component of the system of equations and *θ* is the time constant for the changing, non-stationary frequency. For this EKF, the sensor matrix is 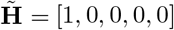 across all time steps *k*. The estimated observation is then

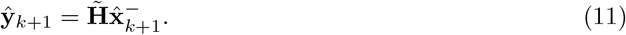

The system of nonlinear ordinary differential equations is solved numerically using Diffrax (Kidger 2022), a JAX ODE solver package in Python. JAX is a scientific computing framework which combines auto-differentiation and TensorFlow’s Accelerated Linear Algebra (XLA). We use the Euler solver with a constant step size.

In this study, we estimate the correlations between the phase estimates of the AOSE and the filter-Hilbert method by computing the mutual information (MI) between them. We estimate a p-value for the MI using a resampling test. The resampling test’s null hypothesis, associated with the chosen permutation type, is that observations within each sample (AOSE phase, filter-Hilbert phase) tuple are drawn from the same underlying distribution and pairings with elements of other samples are assigned at random.

### 2.4. Truth model test of the sequential state estimator

A sequential state estimator, like the EKF of the Adaptive Oscillation State Estimator, must be evaluated to determine whether estimates and error covariance are correct for a given system. While the error and covariance of an estimator will not, in general, equal zero, the estimator can be tested for dynamic consistency. An estimator is defined to be consistent when the state estimate, 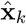, converges in a probabilistic sense to the true state, **x**_*k*_, for large sample sizes (Crassidis and Junkins 2011; Simon 2006). The three criteria of consistency are

1. an unbiased error: E[*e*_*x,k*_] = 0 for all time steps *k*,
2. the true error matches the estimator covariance: 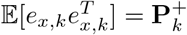, and
3. the errors are white Gaussian distributed: 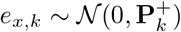,

where 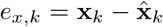. For this study, we will evaluate whether the AOSE is a consistent estimator of, specifically, non-stationary oscillations with variable amplitude and frequency. We use the Normalized Estimation Error Squared (NEES) statistic (Crassidis and Junkins 2011; Simon 2006) to test for estimator consistency:

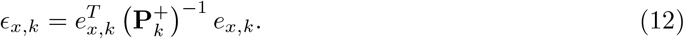

If criteria (1)-(3) for estimator consistency are satisfied, then this implies that the NEES statistic follows a chi-square distribution with *n* degrees of freedom, 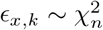 for all time steps *k*, where *n* is the number of state variables tested by NEES. To determine whether NEES is chi-square distributed, we compute an ensemble of NEES values. These NEES values are computed from **truth model testing** across *N* = 1000 Monte Carlo simulations. The average NEES value, 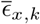, is computed across all simulations; if criteria (1)-(3) are satisfied, then this implies that 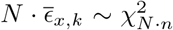, where *N* = 1000 is the number of simulations and *n* is the number of state variables tested. Each simulation runs a ground truth model of a non-stationary oscillation, with varying amplitude and frequency.

The ground truth model is an extended Stuart-Landau model simulated as a system of stochastic differential equations

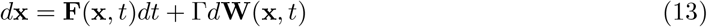

where **W** denotes a Wiener process, Γ ∈ ℝ^5*×*5^ is a process noise covariance matrix

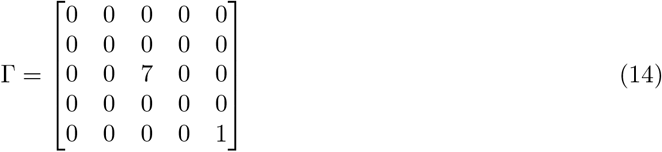

and

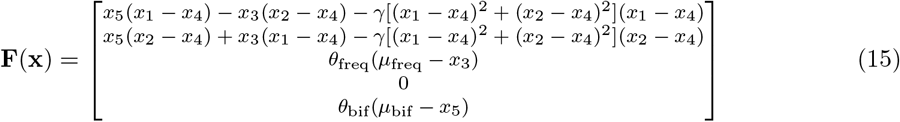

Entries of Γ and **F**(**x**) were chosen such that the frequency and bifurcation parameter of the ground truth SL model would be Ornstein Uhlenbeck processes, making the oscillation frequency and amplitude dynamic.

For each Monte Carlo simulation, the initial condition of the ground truth SL model is drawn from a multivariate normal distribution **x**_0_∼ *N* (**m**_0_, **P**_0_), where 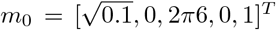 is the initial state of the AOSE filter and

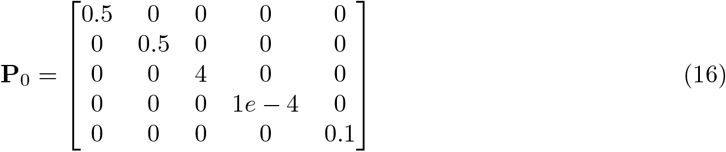

is the initial estimation error covariance matrix of the AOSE filter. The frequency entry of the estimation error covariance matrix is the largest matrix entry, following the assumption that the frequency of a *potential* underlying neural oscillator will be uncertain within a frequency band (e.g., the theta band [4-8] Hz). We test whether the average NEES, 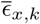, is drawn from a chi-square distribution with *n* degrees of freedom with significance level, *α* = 0.05.

### 2.5. Adaptive estimation of an oscillator’s stimulus response curve using Gaussian process regression

To estimate the phase response curve (PRC) of the Jansen-Rit (JR) model and the amplitude response curve (ARC) from the tremor accelerometry dataset (Cagnan et al. 2017), we use a Gaussian Process (GP) model

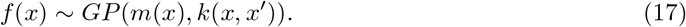

in the BoTorch package in Python (Balandat et al. 2020). A GP is completely specified by a mean function *m* : [0, 2*π*] → R and a covariance function *k* : [0, 2*π*] *×* [0, 2*π*] → R of the process *f* (*x*) (i.e., the underlying PRC). The mean function determines the expected value of *ϕ* = *f* (*x*) at any location *x*

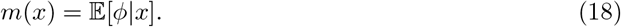

The covariance function determines how deviations from the mean are distributed. If we define *ϕ*^*′*^ = *f* (*x*^*′*^) and *m*^*′*^ = *m*(*x*^*′*^), then we have

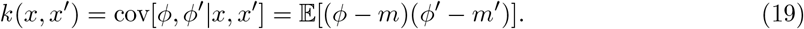

When estimating the JR model’s PRC (Methods 2.1 and Results 3.3), we define the true PRC as the mean function of a GP estimated from a collected dataset of exact phase shifts with no additive noise. Here, we will describe how we estimate this true JR PRC from direct, noise-less observations.

Suppose we observed a function *f* at some set of locations **x∈** ℝ^*n*^, finding the function values ***ϕ*** = *f* (**x**). Let *D* = (**x, *ϕ***) represent the dataset of collected values. We assume the vector of observations is jointly distributed as a Gaussian distribution with any other set of function values from *f* (*x*) ∼ GP (m(*x*), *k*(*x, x*^*′*^)), from a **Gaussian process prior**. The marginal distribution of ***ϕ*** is a Gaussian

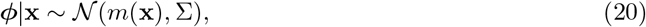

where Σ = *k*(**x, x**) is the Gram matrix formed by evaluating the covariance function for each pair of points: Σ_*ij*_ = *k*(**x**_*i*_, **x**_*j*_). The cross-covariance between any function value *ϕ* = *f* (*x*) and the observation vector ***ϕ*** is given by the covariance function

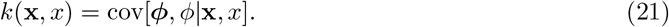

We condition the joint distribution of observations in BoTorch, resulting in a **Gaussian process posterior** on *f*

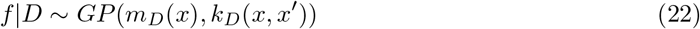

where the updated mean and covariance functions are

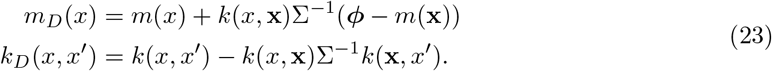

We then define *m*_*D*_(*x*) as the true JR PRC.

In this study, we also add white Gaussian noise to the estimated phase shifts to represent random estimation errors (Results 3.3). To do this, suppose instead that we make observations of the PRC, *f*, at locations **x∈** ℝ^*n*^, measuring noisy phase shifts *y* = ***ϕ*** + ***η***. Suppose estimation errors, ***η*∈** ℝ^*n*^, follow a standard normal distribution

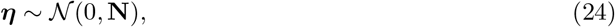

and we collect the measurements into a dataset *D* = (**x, y**). We assume independent, homoskedastic noise with standard deviation *σ* such that the measurement noise covariance matrix is

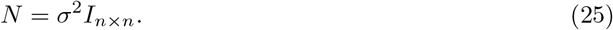

where *I*_*n×n*_ ∈ ℝ^*n×n*^ is the identity matrix. The signal-to-noise ratio of the PRC, *f*, is defined as SNR = *µ*^2^*/σ*^2^, where *µ* is the true PRC and *σ*^2^ is the variance of the additive white Gaussian noise that ”corrupts” the true signal. The prior distribution of the observations from the system is a multivariate normal

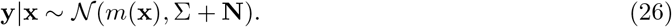

Because the additive noise is independent and identically distributed (i.i.d), the cross-covariance is the same as from the exact, noise-less observation case

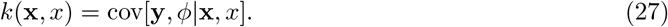

Conditioning on the observed, noise-corrupted values yields a GP posterior with

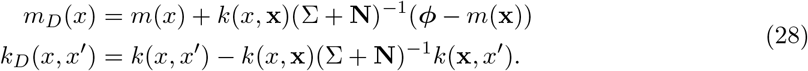

We use the SingleTaskGP class to define a GP in BoTorch with constant prior mean function and radial basis function prior kernel for the prior GP

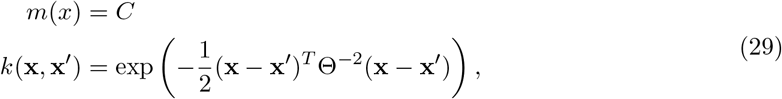

where C is a learned constant and Θ is the kernel length scale parameter. The hyperparameters of the mean and kernel functions of the GP are updated by minimizing the marginal log likelihood objective. Confidence intervals are estimated from the GP posterior. For selecting phases to stimulate the Jansen-Rit model oscillator, we randomly selected a phase using a Sobol sampler, and one phase is sampled every step by the Sobol sampler.

With the scikit-learn package in Python, we compute the *R*^2^ score between the offline PRC estimate (Methods 2.2) and the mean function of the final posterior GP to quantify the degree of similarity between the offline and online PRC estimates (Results 3.4).

## 3 Results

In sections 3.1 and 3.2, we assessed the ability of our Adaptive Oscillation State Estimator (AOSE) to track non-stationary oscillations from a ground truth model using the Stuart-Landau model (section 3.1) in Monte Carlo simulations and from essential tremor accelerometry data from a previously published study (Cagnan et al. 2017) (section 3.2). In the second half, we validated the full Adaptive Oscillation Response Estimation (AORE) pipeline on both simulated and empirical data. First, we demonstrated that the AORE can recover a phase response curve (PRC) from the Jansen-Rit model when the stimulus-induced phase shifts are estimated with random errors (section 3.3). Second, we estimated an amplitude response curve (ARC) from a deep brain stimulation (DBS) experiment (Cagnan et al. 2017) using AOSE. We then compared this online estimated ARC with an ARC estimated from an offline framework, consisting of a bandpass filter, Hilbert Transform, and ordinary linear regression (section 3.4).

### 3.1 Ground truth test of the Adaptive Oscillation State Estimator

In this ground truth model test, the Adaptive Oscillation State Estimator samples observations from the first state variable of the non-stationary Stuart Landau model (Methods 2.3) to recover the underlying observed oscillator state and the hidden oscillator state.

In this *in silico* experiment, we introduced Gaussian white noise to the first state of the ground truth SL model (variance of 0.09) (Figure 3a) The ground truth SL model was driven by Ornstein Uhlenbeck processes in its frequency and bifurcation free-parameter, which yield stochastically variable frequency and amplitude (Figure 3e and 3f). Each time step, a sample from this noise-corrupted observation is used in the AOSE’s measurement correction step to update the oscillation state estimate, the estimation error covariance matrix, and the Kalman gain. Visually, the AOSE recovers the ground truth state variables needed to estimate the phase, frequency, and amplitude envelope of the SL oscillator (Figure 3). In this example, the AOSE filter has mean zero estimation errors that consistently remain within two standard deviations of the estimation error covariance matrix, **P**_*k*_ (Figure 4). This example provides evidence that the AOSE, with chosen process noise covariance matrix, **Q**, and measurement noise covariance matrix, **R**, could be a *consistent* estimator. An estimator is defined to be consistent when the state estimate, 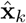, converges in a probabilistic sense to the true state, **x**_*k*_, for large sample sizes (Crassidis and Junkins 2011). To test for filter consistency, we used the Normalized Estimation Error Squared (NEES) statistical test (Crassidis and Junkins 2011; Simon 2006) on 1000 Monte Carlo simulations of the SL model (Figure 5). The observed and hidden states of the SL model were included in the NEES test, including these states’ corresponding block matrix within **P**_*k*_. The initial state of the underlying ground truth SL model was drawn from a multivariate normal distribution, with mean initial condition 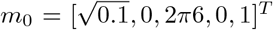 and a covariance matrix equal to the AOSE filter’s estimation error covariance matrix.

**Figure 3:**
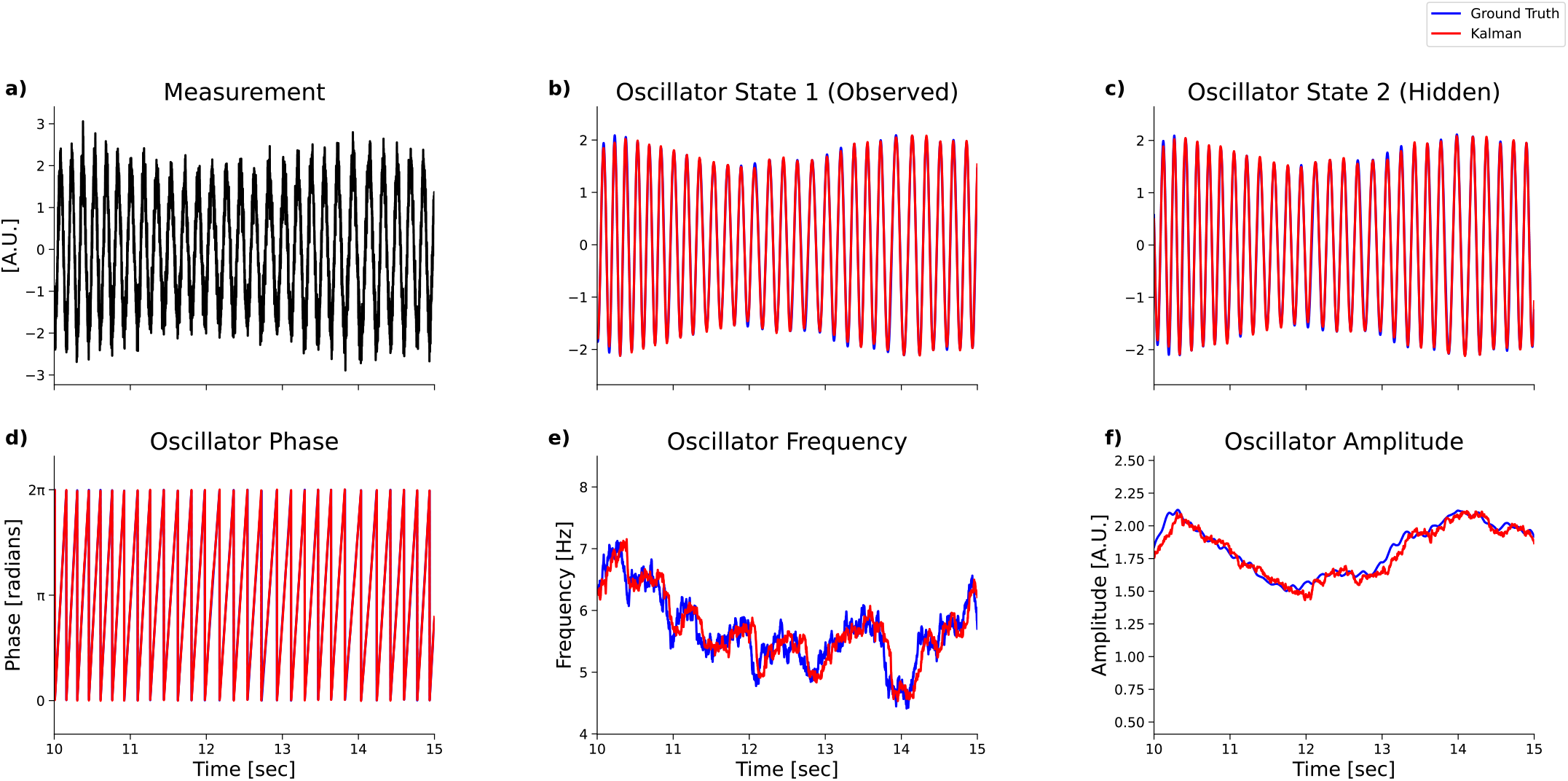
The Adaptive Oscillation State Estimator (AOSE) uses sampled measurements from the first state variable of the non-stationary Stuart Landau model to recover the ground truth observed oscillator state, the hidden oscillator state, and the oscillator frequency state. The ground truth states are in blue and AOSE state estimates are in red. (a) The first SL state variable from the ground truth model, with additional additive white Gaussian noise (variance 0.09), is input into the AOSE, shown in black. (b) The first state variable of the SL model that is observed by the filter. (c) The second state variable of the SL. The second state of the ground truth model is hidden from the AOSE filter. (d) SL phase computed as the inverse tangent of SL states 1 and 2. The estimated phase (red) and true phase (blue) are difficult to distinguish because the estimate agrees closely with ground truth. (e) The third SL state variable, the oscillator frequency, which varies in time as an Ornstein Uhlenbeck process about a mean of 6 Hz. (f) The amplitude envelope of the SL oscillations.

**Figure 4:**
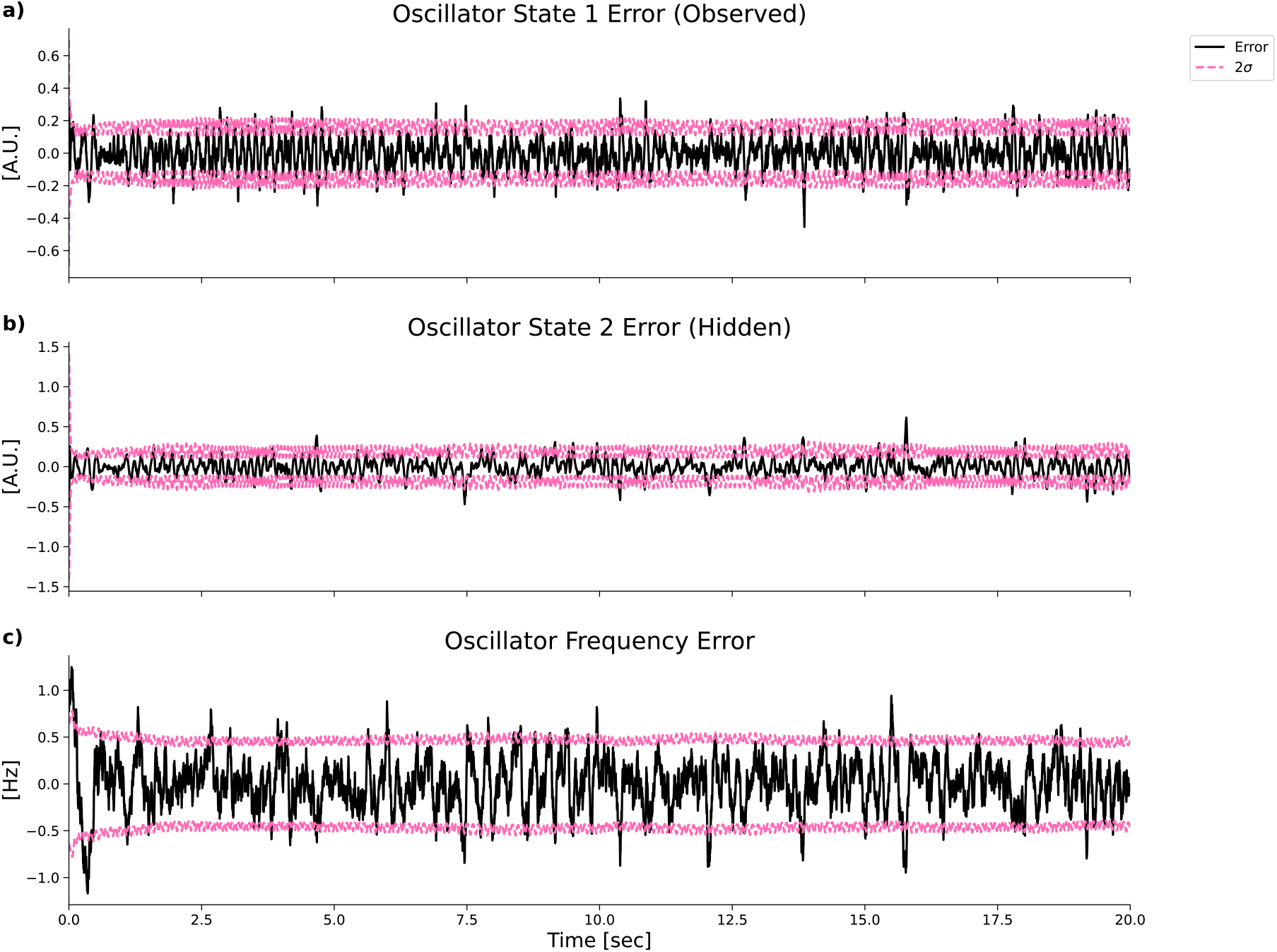
Estimation Errors between the ground truth Stuart Landau model and Adaptive Oscillation State Estimator from the ground truth test (Figure 3). The errors are centered about zero and remain largely within the 2*σ* bounds of the AOSE filter’s estimation error covariance matrix, implying that this is potentially a consistent filter. Errors are in bold black and 2*σ* estimation error bounds are in dashed hot pink. (a) State errors for the first SL state that is indirectly observed by the AOSE filter. (b) State errors for the second SL state that is hidden from the AOSE filter. (c) State errors for the third SL state, the time-varying frequency of the oscillator.

**Figure 5:**
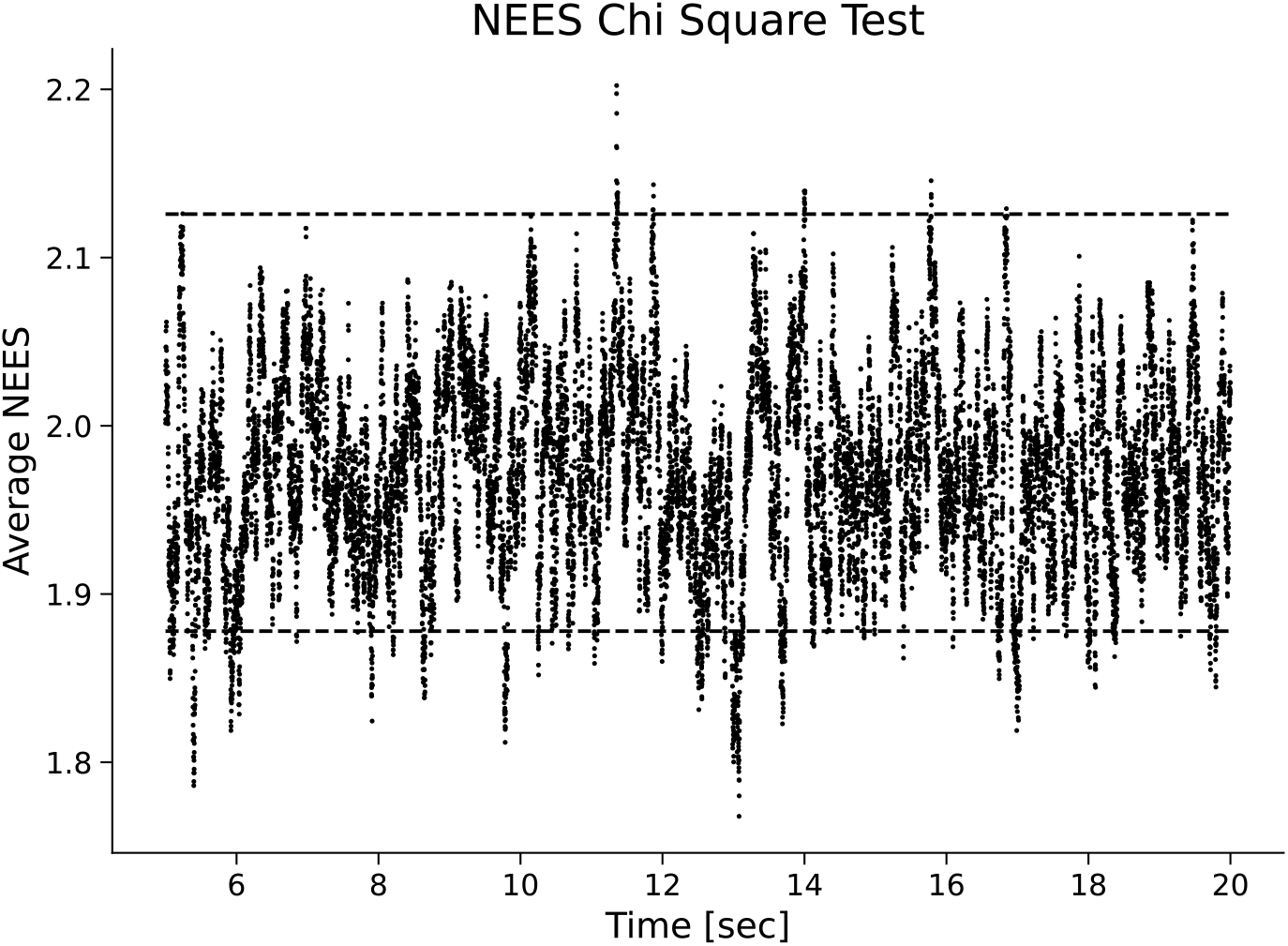
The Adaptive Oscillation State Estimator passes a filter consistency check. Normalized Estimation Error Squared (NEES) test averaged over 1000 Monte Carlo runs in which ground truth Stuart Landau model was initialized from a normal distribution. 95.13% of NEES values lie within 0.05 bounds of a chi square distribution for the Adaptive Oscillation State Estimator. The mean NEES at each time point is a black dot. The 95% chi square significance bounds are dashed lines. These chi square bounds depend on the number of Monte Carlo runs, the number of model states tested, and the target statistical significance.

Over 1000 twenty-second simulations, the AOSE estimation errors were unbiased and the true state error covariance matched the AOSE filter’s estimation error covariance (NEES: 95.13%, *α* = 0.05) (Figure 5) The Adaptive Oscillation State Estimator can recover the underlying ground truth state without underestimating or overestimating the estimation uncertainty, making it a consistent estimator of non-stationary oscillations with, specifically, varying frequency and amplitude.

### 3.2. Adaptive Oscillation State Estimator phase and amplitude match offline estimates from physiologic data

Neural oscillations are typically processed with a bandpass filter and the Hilbert transform (the so called filter-Hilbert method), which has been a useful approach for delivering phase-locked stimulation in the nervous system (Zrenner et al. 2020; Cagnan et al. 2017; Schatza et al. 2022; McNamara, Rothwell, and Sharott 2022). Despite the limitations of the filter-Hilbert method, like edge artifacts and neural waveform distortions, it is still the most common method for extracting neural oscillations from time series data. Because our AOSE method estimates the analytic signal from non-stationary oscillations as well, we hypothesized that our AOSE method would yield comparable results to the filter-Hilbert method on physiologic data. Here, we estimated the phase and amplitude envelope of oscillations in tremor signals from research participant 5 in Cagnan et al. 2017. The AOSE and filter-Hilbert output visually similar results, with the filter-Hilbert amplitude and phase lying within the uncertainty bounds of the AOSE’s filter (Figure 6). The AOSE phase estimate uncertainty grows during distortions in the sinusoidal shape of the signal, whereas the filter-Hilbert approach is unable to quantify this uncertainty. Quantitatively, we estimate the correlations between the filter-Hilbert and AOSE phase by computing the Mutual Information (MI) between these estimates. The MI between the AOSE and filter-Hilbert phase for Participant 5 is MI=1.939 (with p-value of 0.0198) after the resampling test (see Methods 2.3).

**Figure 6:**
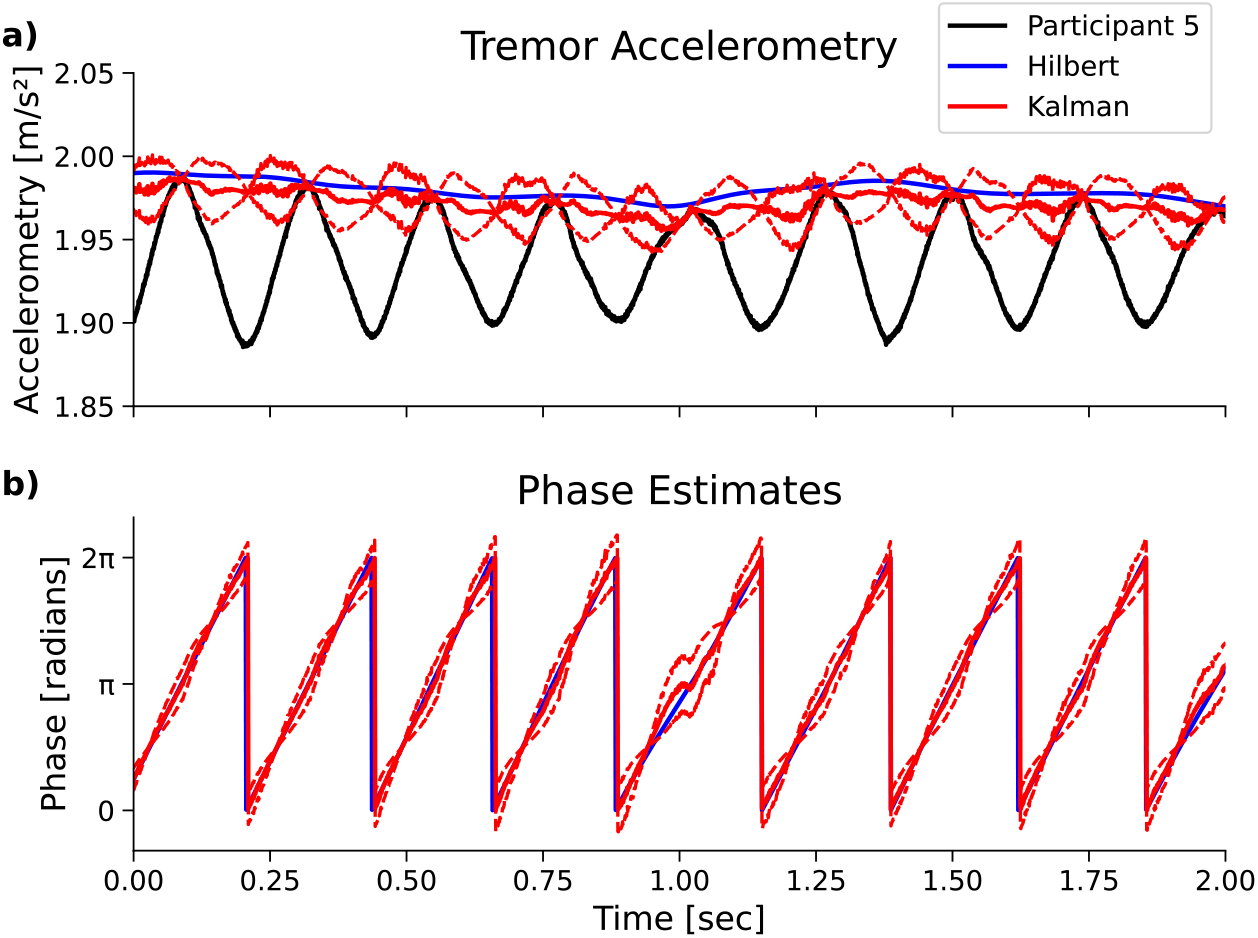
Example of phase and amplitude estimation on 2 seconds of data from Participant 5 in Cagnan et al. 2017 study. (a) Tremor accelerometry from participant in black. The estimated amplitude from applying a band pass filter and Hilbert Transform is in blue. The Kalman filter amplitude estimate is in solid red, with 2 standard deviations about the mean in dashed red. (b) Phase estimate after applying a bandpass filter and Hilbert Transform in solid blue. The Kalman filter phase estimate is in solid red with 2 standard deviations about the mean phase estimate in dashed red. The Hilbert and Kalman phases overlap, especially during large tremor amplitude epochs.

### 3.3. Adaptive Oscillation Response Estimation extracts a similar PRC across different response estimation error levels

Dynamical systems with stable limit cycles are present in many biological systems, including the cardiac pacemaker cells (Mandla, Jung, and Vedantham 2021), the crustacean stomatogastric ganglion’s central pattern generators (Marder and Bucher 2001), and the circadian pacemakers of the suprachiasmatic nucleus of the hypothalamus (Mohawk and Takahashi 2011). The limit cycles of these systems are inherently nonlinear and cannot occur in linear systems (Strogatz 2014). These complex nonlinear oscillators can be reduced to one-dimensional phase models with a phase response curve (PRC), which quantifies the effect of an external perturbation on the phase of a limit cycle (Ermentrout and Terman 2010). Therefore, in this section, we simulate a nonlinear model, the JR model (Jansen and Rit 1995), which is capable of generating self-sustained oscillations and has an inherent PRC.

Our goal is to estimate the JR model’s PRC by perturbing the neural oscillator model and recording stimulus-induced phase shifts via the direct method (Monga et al. 2019). This is analogous to estimating a PRC experimentally from a population of neurons. Behavioral factors such as learning and attention can change the operating point, or state, of a population of neurons (Huang 2021). A change in state of a neural population can alter the dynamic properties of the system, such as the population’s response to external input. If the population is in an oscillatory regime, this change in state could therefore modify the hypothetical population’s PRC. An experimentally estimated PRC should, therefore, be updated over time to track potential changes in the oscillatory state of the system. In this section, we validate a method for updating PRCs sequentially as phase shift samples accumulate over an experiment. For this, we use our Adaptive Oscillation Response Estimation (AORE) framework to estimate a PRC at every sampling step with the direct method for PRC extraction.

Estimating phase shifts (or phase slips) from noisy biological data is a difficult time series analysis task that can be prone to estimation errors. These phase shift estimation errors could bias an experimentally estimated PRC away from the biological system’s true PRC. As a simple model of these estimation errors, we add different magnitudes of white Gaussian noise to the phase shifts extracted from the JR oscillator (Figure 7). The signal-to-noise ratio of the PRC is defined as SNR = *µ*^2^*/σ*^2^, where *µ* is the true PRC and *σ*^2^ is the variance of the additive white Gaussian noise that ”corrupts” the true signal. This white noise can represent random errors in our estimation of stimulation-induced phase shifts along the oscillator’s limit cycle.

**Figure 7:**
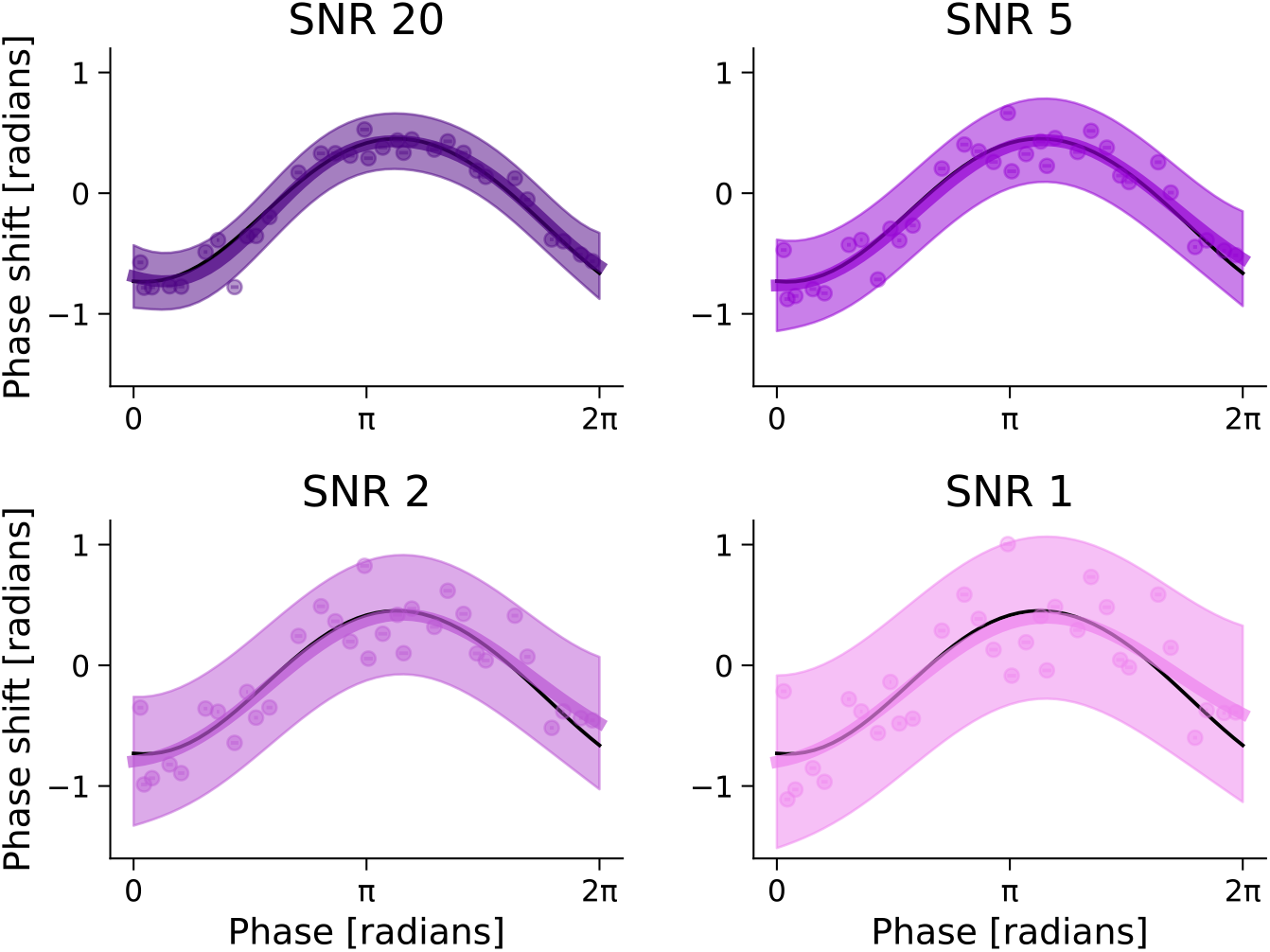
The Gaussian Process PRC estimates, for various phase response estimation error levels, converge to the ideal PRC estimate. Gaussian Process estimates of Stochastic Jansen-Rit model PRC using Adaptive Oscillation Response Estimation. The mean estimate from the Gaussian Process is in bold, and the confidence intervals (2 standard deviations) from the posterior covariance is the filled region. Phase shift samples from stimulating the JR model are light dots. The ideal PRC estimate, with zero additive Gaussian noise applied to phase shifts, is a black solid line. The signal-to-noise ratio (SNR) of the phase-shift synthetic data is modified by additive white Gaussian noise, with variance determined by the selected SNR (SNR = *µ*^2^/*σ*^2^). Top left: SNR=20 (indigo). Top right: SNR=5 (dark violet). Bottom left: SNR=2 (medium orchid). Bottom right: SNR=1 (violet).

Within our AORE method, we use a Gaussian process (GP) model with a constant mean function and radial basis function (RBF) kernel to represent our mean estimate of the PRC and the variance in the phase shifts. Every sampling step, as the new phase data is added to the GP model’s training data, the hyperparameters of the GP are fit to the training data by minimizing the marginal log likelihood objective. After 30 samples, the mean PRC of the GP model from AORE converges to an estimate near what we refer to as the ”ideal” Jansen-Rit PRC (Figures 7 and 8), ”ideal” in the sense that no additive white Gaussian noise is added to the phase shifts. At lower phase shift SNRs, there tends to be a larger deviation between the estimated and ideal PRCs at the boundaries (0 and 2*π* radians) (Figure 7) with AOSE. We averaged over 20 different PRC estimation simulations, and found that the root-mean square error (RMSE) between the AORE PRC estimate and the ideal Jansen-Rit PRC decreases with sampling steps on average across different phase shift SNR levels (Figure 8). Notably, at SNR 1, the RMSE increases before decreasing after sampling step 9. Therefore, PRC estimation error is not guaranteed to monotonically decrease, but as the number of samples increases, the error converges on average after 20 samples, across phase shift SNRs.

**Figure 8:**
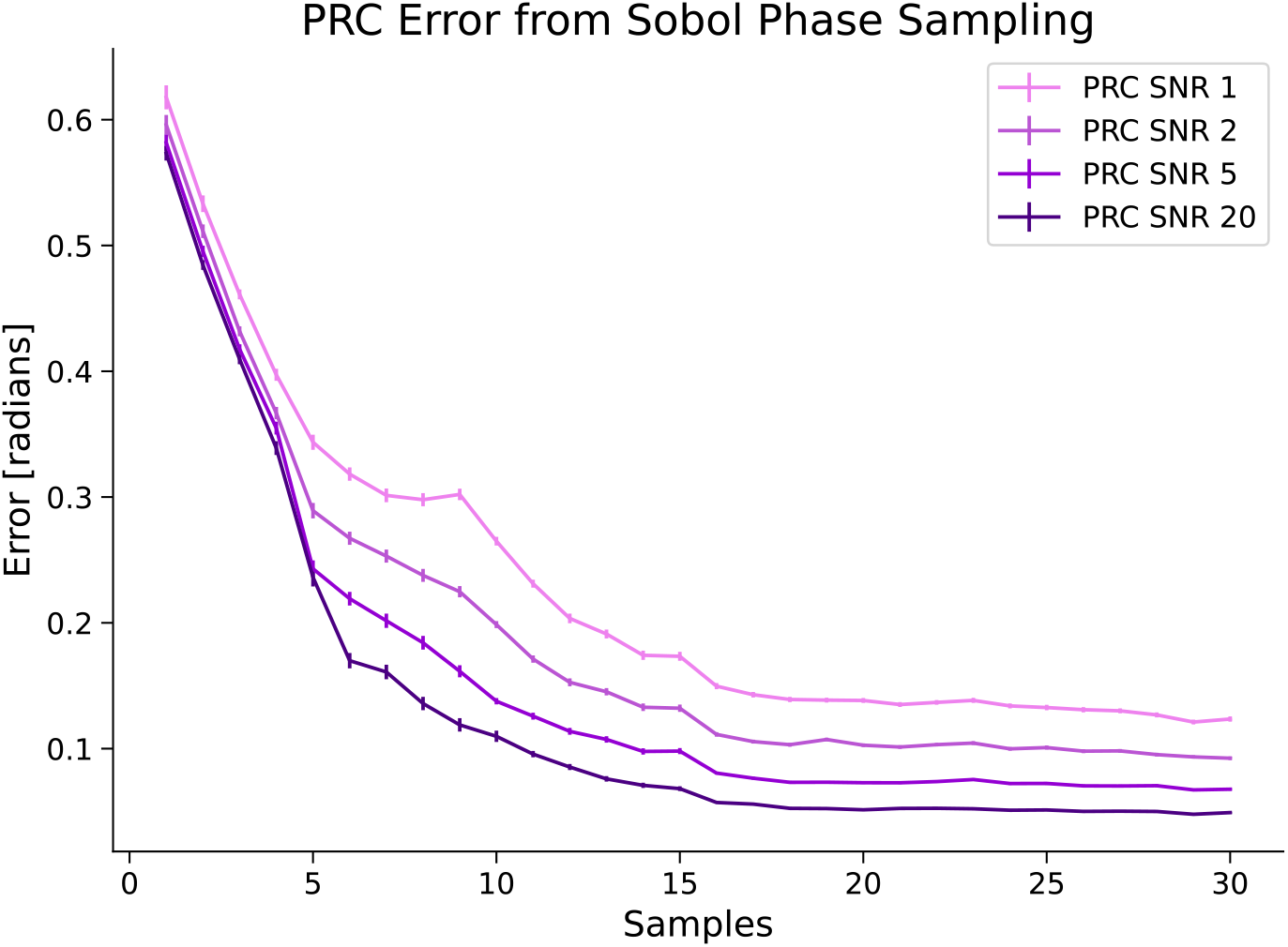
PRC estimation error decreases with sampling steps. The average root-mean square error (RMSE) of Adaptive Oscillation Response Estimation across sampling steps, averaged across 20 simulations. Each sampling step consists of 1 sampled phase and phase shift generated by delivering a phase-locked monophasic pulse to the JR model. The average RMSE is the bold line. The error bars are the standard error of the mean (SEM).

### 3.4. Adaptive Oscillation Response Estimation extracts similar trends from physiologic phasic response data compared to offline analysis

Cagnan and colleagues (Cagnan et al. 2017) delivered deep brain stimulation to the ventral lateral thalamus (VL) that was phase-locked to hand tremor oscillations from ET research participants. They found that stimulating the thalamus at certain tremor phases resulted in significant changes to the amplitude of the tremor rhythms. Another result of this study was that the phase dependence of tremor changes was not uniform across participants (see Methods 2.2), resulting in an individual amplitude response curve (ARC). As an application, we tested whether our Adaptive Oscillation Response Estimator, with sequential oscillatory state estimation and Gaussian process regression, can estimate a similar phase-dependent ARC across participants from Cagnan et al. 2017. For comparison, we estimated the ARCs using an offline analysis pipeline based on the Cagnan study (Methods 2.2). Both the offline analysis and AORE find similar trends in the research participants’ phase-dependent features (Figures 9 and 10). For example, the offline analysis and AORE mean predict negative deflections in the ARC around *π* for Participant 1, a uniform ARC across phase for Participant 3, and a similar positive peak in the ARC for Participant 5. The offline and online ARC estimates were more similar for some participants than others. We quantified the similarity between the offline and online estimates using the *R*^2^ score (Participant 1: *R*^2^ = 0.54, Participant 4L: *R*^2^ = 0.38, Participant 5: *R*^2^ = 0.86, Participant 6: *R*^2^ = 0.26; Figure 10). Compared to the offline ARC, the online ARC from AORE tends to underestimate the phase-dependent changes in tremor oscillatory amplitude induced by DBS. The original study also estimated discrete ARCs across phase space. For ease of comparison to Cagnan et al. 2017, we estimated the midpoint Riemann approximation of the continuous offline ARC and the mean ARC prediction from AORE (Figure 10).

**Figure 9:**
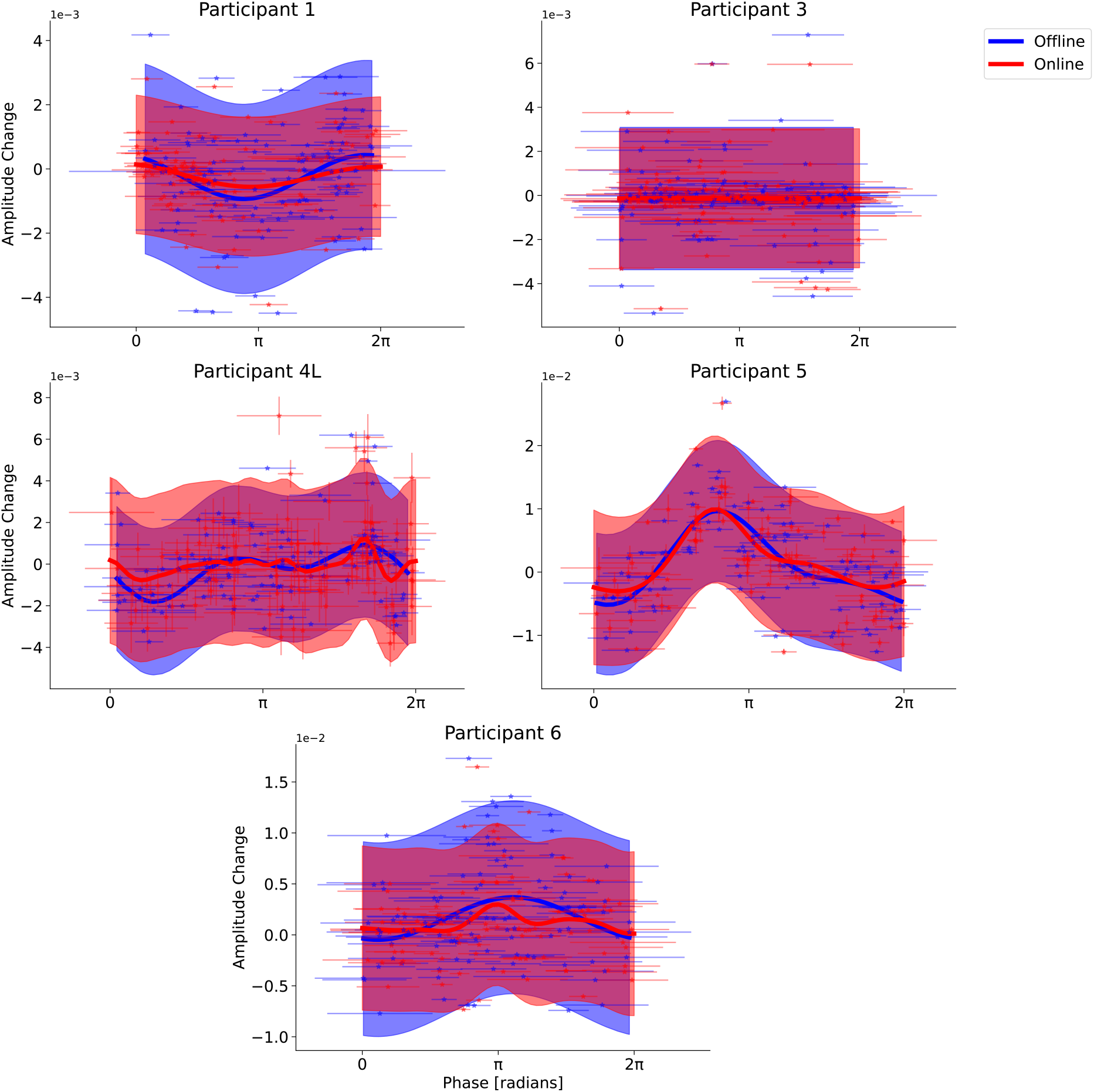
Continuous phase-amplitude response estimates from Adaptive Oscillatory Response Estimation (red) and offline PRC estimation (blue) follow similar trends across stimulated phase. The offline estimate was computed using features extracted from a filter-Hilbert estimate of phase and amplitude and fit with linear regression. Horizontal bars represent the circular average targeted tremor phase in each stimulation block. For the online estimate, vertical bars represent the standard deviation of the Kalman filter’s state estimate (red).

**Figure 10:**
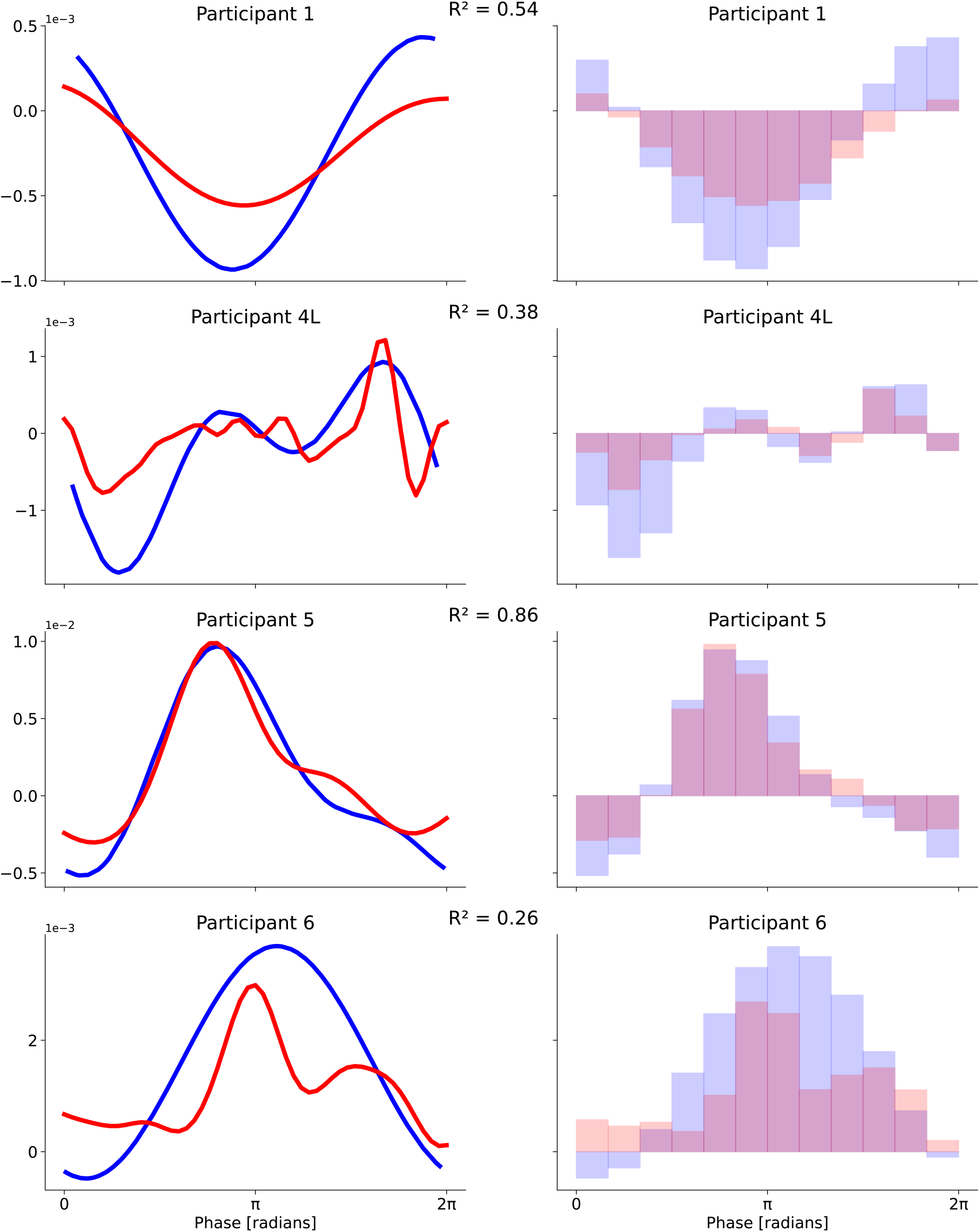
Discrete amplitude response estimates from Adaptive Oscillatory Response Estimation (red) and offline ARC estimation (blue) follow similar trends across stimulated phase. The offline estimate was computed using features extracted from a filter-Hilbert estimate of phase and amplitude and fit with linear regression. Left column: Continuous ARC estimates from the offline analysis (blue) and online analysis with Adaptive Oscillation Response Estimation (red). The *R*^2^ score between the continuous offline and online ARCs is in the upper right hand corner for each participant. Right column: Discrete ARC estimates from offline (light blue) and online analysis (light red) derived from the continuous estimates on the left column using a midpoint Riemann approximation.

## 4 Discussion

In this study, we consistently estimated the state of non-stationary oscillations online from a computational model and from physiologic data (Results 3.1 and 3.2), adaptively estimated the phase response curve (PRC) from a dynamic system in an oscillatory regime (Jansen and Rit 1995) (Results 3.3), and demonstrated adaptive amplitude response curve (ARC) estimation from a previous phase-locked DBS experiment (Cagnan et al. 2017) by sequentially streaming the collected data (Results 3.4). We validated the Adaptive Oscillation State Estimation (AOSE) and Adaptive Oscillation Response Estimation (AORE) frameworks by comparing their performance to either ground truth or an offline estimate. We found that with these frameworks, online estimation of an oscillator’s state and PRC is feasible, even when an oscillator’s amplitude and frequency are time-dependent and the phase-dependent feature is estimated with random errors.

With an adaptive oscillation state and response estimation framework, non-stationary neural oscillations (an observed feature in neural data) can be tracked in a recorded signal. With this framework, applying phase-locked stimulation to this tracked oscillation can yield a PRC if the oscillations originate from a nonlinear oscillator. By discovering the PRC through Bayesian inference, both the mean PRC and uncertainty bounds on the PRC estimate are computed online at each phase response sampling step, which is useful if the form of the PRC is unknown *a priori* and if the PRC is needed to inform an online control policy to modify the dynamics of the neural population (see Extension 5.2). Since some methods to estimate features from neural oscillations assume stationarity (Donoghue, Schaworonkow, and Voytek 2022), this estimation approach will allow the dynamical regime of different areas of the brain to be investigated with less bias introduced by the analysis method. Is a certain population of neurons in the brain behaving dynamically as a nonlinear oscillator with a limit cycle? If so, which type of oscillator? If not, perhaps that population can instead be better represented as a dynamic system with quasi-cycles (Wallace et al. 2011). For instance, a neural population might be in a non-oscillatory state if there is no significant non-uniform trend in the PRC. Further, the dynamic state of neural populations can change with time (Huang 2021), so it will be beneficial to utilize online methods which do not assume that the dynamic state of the brain is static. These adaptive state and response estimator methods could be built upon to help neuroscientists investigate the dynamic regime of different brain regions and bridge the oscillatory features observed in neural data to physiological mechanisms.

There are limitations to the adaptive estimation framework presented in this study, limitations that will need to be addressed before these methods can be applied to other neurophysiological data modalities, like LFP and EEG. (1) The AOSE does not enforce frequency-band constraints. Neural signals are commonly categorized into canonical frequency bands, like alpha and beta, across neuroscience studies. While broadband spectral features are also of interest to the field, if a neuroscientist has a hypothesis based on a particular band, then AOSE in its default form cannot be used. Either data would need to be pre-filtered (adding potential distortions) or the model would need to be extended to emphasize a particular frequency band (see Extension 5.3). (2) The EKF of the AOSE assumes that the measurements it receives contain additive white Gaussian noise, while most biological time series data contains 1*/f* -like pink noise (see Extension 5.5). (3) The AOSE does not simultaneously estimate neural oscillations in multiple frequency bands (see Extension 5.4). (4) Non-sinusoidal waveforms are present in neural oscillations, such as the mu-rhythms of the somatosensory cortex. Currently, the AOSE assumes the oscillator has a circular limit cycle, which implies that the oscillations are assumed to be sinusoidal. As discussed in Results 3.2, the AOSE phase estimate uncertainty grows during distortions in the sinusoidal shape of the signal, whereas the filter-Hilbert approach is unable to quantify this uncertainty (Figure 6). This feature of the AOSE may be useful in neuroscience research, as deviations in the sinusoidality of neural oscillations can create artifacts in adjacent frequency bands of the power spectrum (Schaworonkow 2023; Donoghue, Schaworonkow, and Voytek 2022), leading to the false positive detection of neural oscillations. However, to track non-sinusoidal oscillations, a different nonlinear oscillator model would need to be implemented within AOSE. For non-sinusoidal oscillations to be modeled in the non-stationary oscillation filter, more general limit cycles would need to be modeled, such as ellipses. (5) PRCs and ARCs are defined with an independent phase variable that is defined on a domain with cyclic boundaries. For example, if phase *ϕ* is defined in the domain [0, 2*π*], then the PRC should have the same value at *ϕ* = 0 and *ϕ* = 2*π* to enforce these cyclic boundary conditions. Further, for the PRC to be differentiable at the boundaries, the derivatives should also be the same. The Gaussian process (GP) in the AORE does not enforce cyclic boundary conditions on the domain of its independent variable, as evident in the PRC estimates from the Jansen-Rit model (Figure 7). The GP would need to use a periodic kernel (see Extension 5.1). (6) The AOSE and AORE’s online PRC estimate quality is variable. When recovering ARCs from participants in Cagnan et al., 2017, small differences in the phase and amplitude estimates between the offline and online pipelines could lead to large differences between the offline and online ARCs (Figure 10). The *R*^2^ score between offline and online ARC estimates in this study varied, from 0.26 to 0.86. This may have been due to the different orders of magnitude of the tremor amplitude changes and different complexities in the ARC shape. As we were analyzing a pre-existing dataset, we were limited by the number of stimulation trials per participant.

## 5 Extensions

### 5.1. Gaussian process regression with cyclic boundary conditions

The phase domain [0, 2*π*] is a cyclic domain, with periodic boundaries conditions. A limitation of the Adaptive Oscillation Response Estimator (with Gaussian process regression) is that it does not guarantee for the process *f* (*x*) ∼ GP (m(*x*), *k*(*x, x*^*′*^)) that *f* (0) /= *f* (2*π*). Therefore, it is possible for *f* (0) ≠ *f* (2*π*), which is especially likely as the signal-to-noise ratio (SNR) decreases for the phase-dependent feature of interest (e.g., phase shifts; see Figure 7). To take this into account, the phase-dependent data could be made 2*π*-periodic by repeating it in a domain [ −2*π*, 0] before and in a domain [*™* 2*π*, 4*π*] after the phase domain. For the GP, we could then use the periodic kernel prior

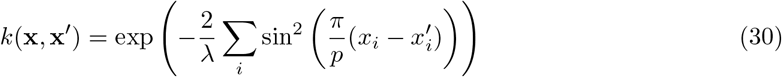

with period *p* = 2*π* and length scale hyperparameter *λ*. This periodic kernel is implemented in the GPyTorch package and is available for use in GP models in the BoTorch package. Because GP kernels can be added and the summation can be itself a GP kernel, this periodic prior kernel could be combined with a RBF kernel (Eq. 25) using the AdditiveKernel class in GPyTorch. This periodic GP could then be fit to the 2*π* extended data and the results outside the domain [0, 2*π*] discarded to guarantee the posterior mean and confidence bounds are 2*π*-periodic.

### 5.2. Gaussian process regression in phasic synchrony controllers

A population of uncoupled model neurons can be desynchronized by stimulating each neuron with an input based on the derivative of the individual neurons’ PRC (Wilson and Moehlis 2014). To extract the PRC derivative from a GP model, one could take the derivative of the GP posterior mean and covariance functions (Eq. 24). Assume the hyperparameters of these functions were fit using a dataset *D*. Then,

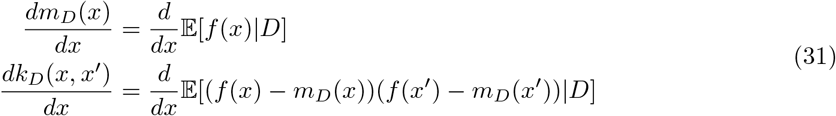

If the kernel is the RBF (Eq. 25), then the function *f* is continuously differentiable, which implies that

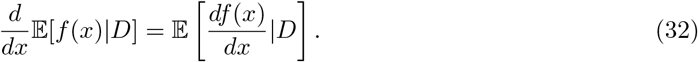

Therefore, the derivative of the mean function *m*(*x*) will equal the expected value of the *derivative* of the process *f* (*x*). One can get a similar expression for the derivative of the covariance function *k*(*x, x*^*′*^). Since the mean and covariance function for the posterior GP are computed each sampling step, then an estimate for the derivative and derivative’s uncertainty can be found for each step that the GP is updated. Therefore, a flexible GP framework should be able to inform neuronal network controllers that utilize the derivative of the PRC (Wilson and Moehlis 2014; Monga et al. 2019), as well, with the added benefit that one can utilize the derivative uncertainty.

### 5.3. Frequency-band constrained oscillation state estimation

Neural oscillations are categorized into specific frequency ranges, such as theta (4-8 Hz), alpha (8-12 Hz), and beta (12-30 Hz), that each show different changes during motor movements, sensory processing, and cognitive tasks (Canolty and Knight 2010). The boundaries of these frequency bands are arbitrary and vary across studies, but they do help compare electrophysiological results across studies. Therefore, it is useful in neuroscience to estimate properties of oscillations, such as phase and amplitude, within these band constraints. With the filter-Hilbert approach, a bandpass filter is applied to electrophysiological time series data, and the boundaries of this bandpass filter are typically chosen to match a frequency band of interest that a researcher hypothesizes is a relevant feature of interest. Our Adaptive Oscillation State Estimator (AOSE) does not currently restrict the frequency state of its internal Stuart-Landau model within a frequency band. However, the method can be extended to accommodate this by using a constrained Kalman Filter.

Based on Simon 2006, suppose that the states of our filter satisfy some imposed inequality constraint **Dx** ≤ **d** such that

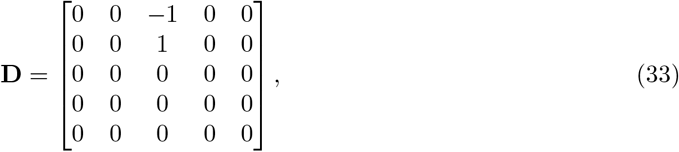

**x** = [*x*_1_, *x*_2_, *x*_3_, *x*_4_, *x*_5_]^*T*^ is the state of the SL model (where *x*_3_ is the oscillator frequency; see Methods) and **d** = [*−*2*πa*, 2*πb*, 0, 0, 0]^*T*^ is the constraint vector. This implies that the frequency is bounded between 2*πa ≤ x*_3_ ≤ 2*πb*, for a frequency band [*a, b*] Hz. We can use the ”projection approach” (Simon 2006) to obtain a constrained state estimate by solving the quadratic programming problem

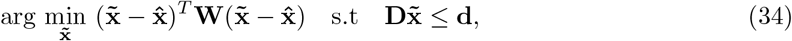

where **W** is any positive definite weight matrix, with matrix elements of **W** chosen based on our confidence in the variables of the unconstrained state estimate from the Kalman filter. Common choices for are 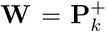 and **W** = *I* (Simon 2006). Since the constraints on the frequency band are known *a priori*, the solution to this quadratic programming problem is also the solution to the equality-constrained problem

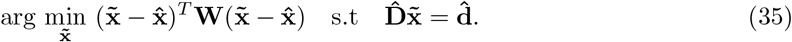

with

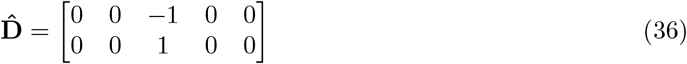

and 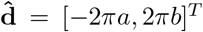, which only uses the rows of **D** and elements of **d** corresponding to our frequency constraints. The solution to the equality-constrained problem is (Simon 2006)

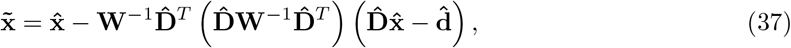

which is just a linear system of equations. Therefore, the frequency-constrained state estimate, 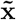, can be derived from the AOSE’s unconstrained oscillator state estimate, 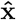, at every time step.

### 5.4. Estimate oscillations in multiple frequency bands with interacting multiple model estimation

Some neuroscience theories posit that LFP and EEG oscillations in different frequency bands, like theta and alpha, represent signals from distinct nonlinear oscillators in spatially separate populations of neurons. In these theories, these neural populations, with limit cycles in different frequency bands, interact as weakly coupled oscillators that influence the phase and amplitude of their rhythms. This provides one possible explanation for the statistical dependence found between the phases (or phase and amplitude) of different frequency bands, termed cross-frequency coupling (CFC), which could coordinate information processing between brain regions. Testing the validity of this theory of cross-frequency coupled oscillators has proven difficult. For example, non-sinusoidal neural waveforms in LFP and EEG, such as the mu-rhythm, are represented as multiple harmonic peaks in different frequency bands of the power spectrum. This can result in the spurious claim that these peaks represents distinct oscillating neural populations that physiologically interact via CFC (Donoghue, Schaworonkow, and Voytek 2022). Simultaneous, model-based estimation of different frequency-band neural oscillators may help researchers avoid these false positives. Currently, the Adaptive Oscillation State Estimator only tracks the frequency of a single oscillator. As discussed in 5.3., this method can be extended to track frequency-constrained oscillations. As a further extension, multiple frequencies could be estimated simultaneously using an interacting multiple model (IMM) estimator. This IMM implementation could weigh each oscillator frequency estimator by the likelihood that the oscillator is present, using the residual between model estimate and measurement (Crassidis and Junkins 2011). Transitions between the different frequency bands would follow a Markov sequence, with transition probabilities between the oscillator models weighted by the frequency likelihoods.

### 5.5. Sequential state estimation with colored measurement noise

Neural recordings contain both oscillatory and non-oscillatory features (Donoghue, Schaworonkow, and Voytek 2022). This aperiodic, 1/f-like activity in neural data could confound and bias the AOSE and other sequential state estimators of neural oscillations. While the Extended Kalman filter of the AOSE assumes measurement noise is additive white Gaussian, one can design a Kalman filter that can optimally handle colored noise. We outline a brief sketch for how this can be done based on previous work (Crassidis and Junkins 2011; Simon 2006). We can pass additive white Gaussian measurement noise through a so-called shaping filter such that

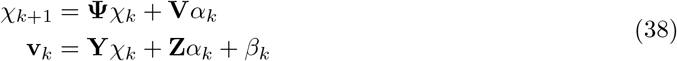

where *α*_*k*_ and *β*_*k*_ are both additive white Gaussian processes with covariances, **Λ** and **Φ**, respectively. Assuming *α*_*k*_ and *β*_*k*_ are uncorrelated, the new measurement noise covariance matrix is given by

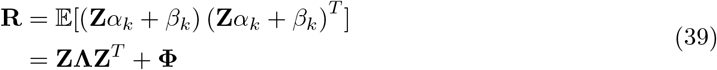

We can then augment the state vector with the shaping filter state during the Kalman filter’s prediction and measurement estimation steps (see Methods 2.3)

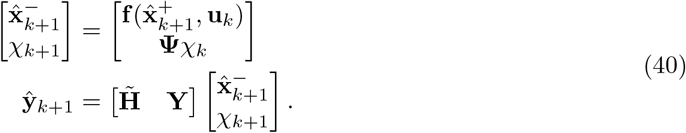

Assuming that the process noise, **w**_*k*_, and *α*_*k*_ are uncorrelated, the new process noise covariance matrix is

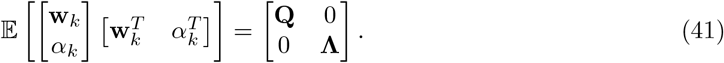

But, for the new augmented system, the new process noise and measurement noise are now correlated, with covariance

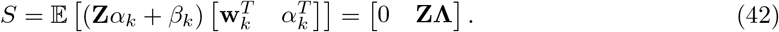

However, our current filter in the AOSE assumes the process and measurement noise are uncorrelated (Methods 2.3). One would need to use a new Kalman filter formulation that can handle correlations between process and measurement noise (Crassidis and Junkins 2011). A linear system of equations, like this shaping filter augmentation to the EKF, may be able to recreate the power spectral scaling of colored noise recorded in cortex (Miller et al. 2009).

## Notes

### Competing Interest Statement

The authors have declared no competing interest.

## References

Abuzaid, Ali H., Ibrahim B. Mohamed, and Abdul G. Hussin (Sept. 2012). “Boxplot for circular variables”. en. In: Computational Statistics 27.3, pp. 381–392. ISSN: 1613-9658. doi: 10.1007/s00180-011-0261-5. URL: 10.1007/s00180-011-0261-5 (visited on 05/28/2024).

Alonso, David, Alan J McKane, and Mercedes Pascual (Dec. 2006). “Stochastic amplification in epidemics”. In: Journal of The Royal Society Interface 4.14. Publisher: Royal Society, pp. 575–582. doi: 10.1098/rsif.2006.0192. URL: https://royalsocietypublishing.org/doi/10.1098/rsif.2006.0192 (visited on 05/09/2024).

Balandat, Maximilian et al. (Dec. 2020). BoTorch: A Framework for Efficient Monte-Carlo Bayesian Optimization. Number: 1910.06403 arXiv:1910.06403 [cs, math, stat]. URL: http://arxiv.org/abs/1910.06403 (visited on 05/26/2023).

Brunel, Nicolas (May 2000). “Dynamics of Sparsely Connected Networks of Excitatory and Inhibitory Spiking Neurons”. en. In: Journal of Computational Neuroscience 8.3, pp. 183–208. ISSN: 1573-6873. doi: 10.1023/A:1008925309027. URL: 10.1023/A:1008925309027 (visited on 03/06/2024).

Cagnan, Hayriye et al. (Jan. 2017). “Stimulating at the right time: phase-specific deep brain stimulation”. In: Brain 140.1, pp. 132–145. ISSN: 0006-8950. doi: 10.1093/brain/aww286. URL: 10.1093/brain/aww286 (visited on 06/13/2022).

Canolty, Ryan T. and Robert T. Knight (Nov. 2010). “The functional role of cross-frequency coupling”. English. In: Trends in Cognitive Sciences 14.11. Publisher: Elsevier, pp. 506–515. ISSN: 1364-6613, 1879-307X. doi: 10.1016/j.tics.2010.09.001. URL: https://www.cell.com/trends/cognitive-sciences/abstract/S1364-6613(10)00206-8 (visited on 05/08/2024).

Connolly, Mark J. et al. (May 2021). “Multi-objective data-driven optimization for improving deep brain stimulation in Parkinson’s disease”. en. In: Journal of Neural Engineering 18.4. Publisher: IOP Publishing, p. 046046. ISSN: 1741-2552. doi: 10.1088/1741-2552/abf8ca. URL: 10.1088/1741-2552/abf8ca (visited on 05/28/2024).

Crassidis, John L. and John L. Junkins (Oct. 2011). Optimal Estimation of Dynamic Systems, Second Edition. en. Google-Books-ID: ITKKkBFxgNsC. CRC Press. ISBN: 978-1-4398-3985-0.

Doelling, Keith B. and M. Florencia Assaneo (May 2021). “Neural oscillations are a start toward understanding brain activity rather than the end”. en. In: PLOS Biology 19.5. Publisher: Public Library of Science, e3001234. ISSN: 1545-7885. doi: 10.1371/journal.pbio.3001234. URL: https://journals.plos.org/plosbiology/article?id=10.1371/journal.pbio.3001234 (visited on 02/06/2023).

Donoghue, Thomas, Natalie Schaworonkow, and Bradley Voytek (2022). “Methodological considerations for studying neural oscillations”. en. In: European Journal of Neuroscience 55.11-12. eprint: https://onlinelibrary.wiley.com/doi/pdf/10.1111/ejn.15361, pp. 3502–3527. ISSN: 1460-9568. doi: 10.1111/ejn.15361. URL: https://onlinelibrary.wiley.com/doi/abs/10.1111/ejn.15361 (visited on 10/04/2022).

Ermentrout, G. Bard and David H. Terman (2010). Mathematical Foundations of Neuroscience. Vol. 35. Interdisciplinary Applied Mathematics. New York, NY: Springer. ISBN: 978-0-387-87707-5 978-0-387-87708-2. doi: 10.1007/978-0-387-87708-2. URL: http://link.springer.com/10.1007/978-0-387-87708-2 (visited on 12/06/2022).

Garnett, Roman (2023). Bayesian Optimization. Cambridge: Cambridge University Press. ISBN: 978-1-108-42578-0. doi: 10.1017/9781108348973. URL: https://www.cambridge.org/core/books/bayesian-optimization/11AED383B208E7F22A4CE1B5BCBADB44 (visited on 06/27/2023).

Gerstner, Wulfram et al. (2014). Neuronal Dynamics: From Single Neurons to Networks and Models of Cognition. Cambridge: Cambridge University Press. ISBN: 978-1-107-06083-8. doi: 10.1017/CBO9781107447615. URL: https://www.cambridge.org/core/books/neuronal-dynamics/75375090046733765596191E23B2959D (visited on 05/28/2024).

Gonçalves, Pedro J et al. (Sept. 2020). “Training deep neural density estimators to identify mechanistic models of neural dynamics”. In: eLife 9. Ed. by John R Huguenard, Timothy O’Leary, and Mark S Goldman. Publisher: eLife Sciences Publications, Ltd, e56261. ISSN: 2050-084X. doi: 10.7554/eLife.56261. URL: 10.7554/eLife.56261 (visited on 05/28/2024).

Grado, Logan L., Matthew D. Johnson, and Theoden I. Netoff (Dec. 2018). “Bayesian adaptive dual control of deep brain stimulation in a computational model of Parkinson’s disease”. en. In: PLOS Computational Biology 14.12. Publisher: Public Library of Science, e1006606. ISSN: 1553-7358. doi: 10.1371/journal.pcbi.1006606. URL: https://journals.plos.org/ploscompbiol/article?id=10.1371/journal.pcbi.1006606 (visited on 06/17/2022).

Gramfort, Alexandre et al. (2013). “MEG and EEG data analysis with MNE-Python”. In: Frontiers in Neuroscience 7. ISSN: 1662-453X. URL: https://www.frontiersin.org/articles/10.3389/fnins.2013.00267 (visited on 06/27/2023).

Grimbert, François and Olivier Faugeras (Dec. 2006). “Bifurcation Analysis of Jansen’s Neural Mass Model”. In: Neural Computation 18.12, pp. 3052–3068. ISSN: 0899-7667. doi: 10.1162/neco.2006.18.12.3052. URL: 10.1162/neco.2006.18.12.3052 (visited on 04/24/2023).

Huang, Chengcheng (Oct. 2021). “Modulation of the dynamical state in cortical network models”. In: Current Opinion in Neurobiology. Computational Neuroscience 70, pp. 43–50. ISSN: 0959-4388. doi: 10.1016/j.conb.2021.07.004. URL: https://www.sciencedirect.com/science/article/pii/S0959438821000738 (visited on 03/20/2024).

Jansen, Ben H. and Vincent G. Rit (Sept. 1995). “Electroencephalogram and visual evoked potential generation in a mathematical model of coupled cortical columns”. en. In: Biological Cybernetics 73.4, pp. 357–366. ISSN: 1432-0770. doi: 10.1007/BF00199471. URL: 10.1007/BF00199471 (visited on 12/06/2022).

Kehnemouyi, Yasmine M et al. (Feb. 2021). “Modulation of beta bursts in subthalamic sensorimotor circuits predicts improvement in bradykinesia”. In: Brain 144.2, pp. 473–486. ISSN: 0006-8950. doi: 10.1093/brain/awaa394. URL: 10.1093/brain/awaa394 (visited on 05/14/2024).

Kidger, Patrick (Feb. 2022). On Neural Differential Equations. Number: 2202.02435 arXiv:2202.02435 [cs, math, stat]. doi: 10.48550/arXiv.2202.02435. URL: http://arxiv.org/abs/2202.02435 (visited on 02/06/2023).

Kim, Myunghee et al. (Sept. 2017). “Human-in-the-loop Bayesian optimization of wearable device parameters”. en. In: PLOS ONE 12.9. Publisher: Public Library of Science, e0184054. ISSN: 1932-6203. doi: 10.1371/journal.pone.0184054. URL: https://journals.plos.org/plosone/article?id=10.1371/journal.pone.0184054 (visited on 05/28/2024).

Kuznetsov, Yuri A. (2004). Elements of Applied Bifurcation Theory. Ed. by S. S. Antman, J. E. Marsden, and L. Sirovich. Vol. 112. Applied Mathematical Sciences. New York, NY: Springer. ISBN: 978-1-4419-1951-9 978-1-4757-3978-7. doi: 10.1007/978-1-4757-3978-7. URL: http://link.springer.com/10.1007/978-1-4757-3978-7 (visited on 06/28/2023).

Mandla, Ravi, Catherine Jung, and Vasanth Vedantham (July 2021). “Transcriptional and Epigenetic Landscape of Cardiac Pacemaker Cells: Insights Into Cellular Specialization in the Sinoatrial Node”. English. In: Frontiers in Physiology 12. Publisher: Frontiers. ISSN: 1664-042X. doi:10.3389/fphys.2021.712666. URL: https://www.frontiersin.org/journals/physiology/articles/10.3389/fphys.2021.712666/full (visited on 05/28/2024).

Marder, Eve and Dirk Bucher (Nov. 2001). “Central pattern generators and the control of rhythmic movements”. In: Current Biology 11.23, R986–R996. ISSN: 0960-9822. doi: 10.1016/S0960-9822(01)00581-4. URL: https://www.sciencedirect.com/science/article/pii/S0960982201005814 (visited on 05/28/2024).

McKane, A. J. and T. J. Newman (June 2005). “Predator-Prey Cycles from Resonant Amplification of Demographic Stochasticity”. In: Physical Review Letters 94.21. Publisher: American Physical Society, p. 218102. doi: 10.1103/PhysRevLett.94.218102. URL: https://link.aps.org/doi/10.1103/PhysRevLett.94.218102 (visited on 05/09/2024).

McNamara, Colin G., Max Rothwell, and Andrew Sharott (Nov. 2022). “Stable, interactive modulation of neuronal oscillations produced through brain-machine equilibrium”. en. In: Cell Reports 41.6, p. 111616. ISSN: 22111247. doi: 10.1016/j.celrep.2022.111616. URL: https://linkinghub.elsevier.com/retrieve/pii/S2211124722014851 (visited on 05/17/2023).

Miller, Kai J. et al. (Dec. 2009). “Power-Law Scaling in the Brain Surface Electric Potential”. en. In: PLOS Computational Biology 5.12. Publisher: Public Library of Science, e1000609. ISSN: 1553-7358. doi: 10.1371/journal.pcbi.1000609. URL: https://journals.plos.org/ploscompbiol/article?id=10.1371/journal.pcbi.1000609 (visited on 05/23/2024).

Mohawk, Jennifer A. and Joseph S. Takahashi (July 2011). “Cell autonomy and synchrony of suprachiasmatic nucleus circadian oscillators”. In: Trends in Neurosciences 34.7, pp. 349–358. ISSN: 0166-2236. doi: 10.1016/j.tins.2011.05.003. URL: https://www.sciencedirect.com/science/article/pii/S0166223611000749 (visited on 05/28/2024).

Monga, Bharat et al. (Apr. 2019). “Phase reduction and phase-based optimal control for biological systems: a tutorial”. en. In: Biological Cybernetics 113.1, pp. 11–46. ISSN: 1432-0770. doi: 10.1007/s00422-018-0780-z. URL: 10.1007/s00422-018-0780-z (visited on 06/08/2022).

Nagrale, Sumedh S. et al. (May 2023). “In silico development and validation of Bayesian methods for optimizing deep brain stimulation to enhance cognitive control”. en. In: Journal of Neural Engineering 20.3. Publisher: IOP Publishing, p. 036015. ISSN: 1741-2552. doi: 10.1088/1741-2552/acd0d5. URL: 10.1088/1741-2552/acd0d5 (visited on 05/28/2024).

Newson, Jennifer J. and Tara C. Thiagarajan (Jan. 2019). “EEG Frequency Bands in Psychiatric Disorders: A Review of Resting State Studies”. English. In: Frontiers in Human Neuroscience 12. Publisher: Frontiers. ISSN: 1662-5161. doi: 10.3389/fnhum.2018.00521. URL: https://www.frontiersin.org/articles/10.3389/fnhum.2018.00521 (visited on 05/14/2024).

Peterson, Erik J. et al. (July 2023). “Aperiodic Neural Activity is a Better Predictor of Schizophrenia than Neural Oscillations”. en. In: Clinical EEG and Neuroscience 54.4. Publisher: SAGE Publications Inc, pp. 434–445. ISSN: 1550-0594. doi: 10.1177/15500594231165589. URL: 10.1177/15500594231165589 (visited on 05/28/2024).

Rasmussen, Carl Edward and Christopher K. I. Williams (Nov. 2005). Gaussian Processes for Machine Learning. en. doi: 10.7551/mitpress/3206.001.0001. URL: https://direct.mit.edu/books/book/2320/Gaussian-Processes-for-Machine-Learning (visited on 12/06/2022).

Renart, Alfonso et al. (Jan. 2010). “The Asynchronous State in Cortical Circuits”. In: Science 327.5965. Publisher: American Association for the Advancement of Science, pp. 587–590. doi: 10.1126/science.1179850. URL: https://www.science.org/doi/10.1126/science.1179850 (visited on 06/12/2023).

Richardson, Magnus J. E. (Nov. 2008). “Spike-train spectra and network response functions for non-linear integrate-and-fire neurons”. en. In: Biological Cybernetics 99.4, pp. 381–392. ISSN: 1432-0770. doi: 10.1007/s00422-008-0244-y. URL: 10.1007/s00422-008-0244-y (visited on 01/26/2024).

Scangos, Katherine Wilson et al. (2021). “Distributed Subnetworks of Depression Defined by Direct Intracranial Neurophysiology”. In: Frontiers in Human Neuroscience 15. ISSN: 1662-5161. URL: https://www.frontiersin.org/articles/10.3389/fnhum.2021.746499 (visited on 09/03/2022).

Schatza, Mark J. et al. (Jan. 2022). “Toolkit for Oscillatory Real-time Tracking and Estimation (TORTE)”. en. In: Journal of Neuroscience Methods 366, p. 109409. ISSN: 0165-0270. doi: 10.1016/j.jneumeth.2021.109409. URL: https://www.sciencedirect.com/science/article/pii/S0165027021003447 (visited on 01/27/2022).

Schaworonkow, Natalie (Sept. 2023). “Overcoming harmonic hurdles: Genuine beta-band rhythms vs. contributions of alpha-band waveform shape”. In: Imaging Neuroscience 1, pp. 1–8. ISSN: 2837-6056. doi: 10.1162/imag_a_00018. URL: 10.1162/imag_a_00018 (visited on 09/13/2023).

Schwalger, Tilo and Anton V Chizhov (Oct. 2019). “Mind the last spike — firing rate models for mesoscopic populations of spiking neurons”. In: Current Opinion in Neurobiology. Computational Neuroscience 58, pp. 155–166. ISSN: 0959-4388. doi: 10.1016/j.conb.2019.08.003. URL:https://www.sciencedirect.com/science/article/pii/S095943881930039X (visited on 03/20/2024).

Shadlen, Michael N. and William T. Newsome (May 1998). “The Variable Discharge of Cortical Neurons: Implications for Connectivity, Computation, and Information Coding”. en. In: Journal of Neuroscience 18.10. Publisher: Society for Neuroscience Section: ARTICLE, pp. 3870–3896. ISSN: 0270-6474, 1529-2401. doi: 10.1523/JNEUROSCI.18-10-03870.1998. URL: https://www.jneurosci.org/content/18/10/3870 (visited on 05/28/2024).

Shupe, Larry and Eberhard Fetz (Mar. 2021). “An Integrate-and-Fire Spiking Neural Network Model Simulating Artificially Induced Cortical Plasticity”. en. In: eNeuro 8.2. Publisher: Society for Neuroscience Section: Research Article: New Research. ISSN: 2373-2822. doi: 10.1523/ENEURO.0333-20.2021. URL: https://www.eneuro.org/content/8/2/ENEURO.0333-20.2021 (visited on 06/12/2023).

Simon, Dan (June 2006). Optimal State Estimation: Kalman, H Infinity, and Nonlinear Approaches. en. Google-Books-ID: UiMVoP 7TZkC. John Wiley & Sons. ISBN: 978-0-470-04533-6.

Strogatz, Steven H. (July 2014). Nonlinear Dynamics and Chaos: With Applications to Physics, Biology, Chemistry, and Engineering, Second Edition. English. 2nd edition. Boca Raton London New York: Westview Press. ISBN: 978-0-8133-4910-7.

Vreeswijk, C. van and H. Sompolinsky (Dec. 1996). “Chaos in Neuronal Networks with Balanced Excitatory and Inhibitory Activity”. In: Science 274.5293. Publisher: American Association for the Advancement of Science, pp. 1724–1726. doi: 10.1126/science.274.5293.1724. URL: https://www.science.org/doi/10.1126/science.274.5293.1724 (visited on 05/28/2024).

Wallace, Edward et al. (May 2011). “Emergent Oscillations in Networks of Stochastic Spiking Neurons”. en. In: PLOS ONE 6.5. Publisher: Public Library of Science, e14804. ISSN: 1932-6203. doi: 10.1371/journal.pone.0014804. URL: https://journals.plos.org/plosone/article?id=10.1371/journal.pone.0014804 (visited on 05/08/2024).

Weerasinghe, Gihan et al. (Aug. 2021). “Optimal closed-loop deep brain stimulation using multiple independently controlled contacts”. en. In: PLOS Computational Biology 17.8. Publisher: Public Library of Science, e1009281. ISSN: 1553-7358. doi: 10.1371/journal.pcbi.1009281. URL: https://journals.plos.org/ploscompbiol/article?id=10.1371/journal.pcbi.1009281 (visited on 03/11/2022).

Wilson, Dan and Jeff Moehlis (Mar. 2014). “Optimal Chaotic Desynchronization for Neural Populations”. en. In: SIAM Journal on Applied Dynamical Systems. Publisher: Society for Industrial and Applied Mathematics. doi: 10.1137/120901702. URL: https://epubs.siam.org/doi/10.1137/120901702 (visited on 12/06/2022).

Wodeyar, Anirudh et al. (Sept. 2021). “A state space modeling approach to real-time phase estimation”. In: eLife 10. Ed. by Frances K Skinner, Laura L Colgin, and Milad Lankarany. Publisher: eLife Sciences Publications, Ltd, e68803. ISSN: 2050-084X. doi: 10.7554/eLife.68803. URL: 10.7554/eLife.68803 (visited on 01/27/2022).

Zanos, Stavros et al. (Aug. 2018). “Phase-Locked Stimulation during Cortical Beta Oscillations Produces Bidirectional Synaptic Plasticity in Awake Monkeys”. eng. In: Current biology: CB 28.16, 2515–2526.e4. ISSN: 1879-0445. doi: 10.1016/j.cub.2018.07.009.

Zrenner, Brigitte et al. (Jan. 2020). “Brain oscillation-synchronized stimulation of the left dorsolateral prefrontal cortex in depression using real-time EEG-triggered TMS”. In: Brain Stimulation 13.1, pp. 197–205. ISSN: 1935-861X. doi: 10.1016/j.brs.2019.10.007. URL: https://www.sciencedirect.com/science/article/pii/S1935861X19304140 (visited on 08/14/2023).

